# Revisiting the African mtDNA Landscape: A Continental Update from Complete Mitochondrial Genomes

**DOI:** 10.1101/2025.04.05.647361

**Authors:** Imke Lankheet, Afifa Chowdhury, Christian Tellgren-Roth, Cécile Jolly, André E. R. Soares, Miguel de Navascués, Sara Pacchiarotti, Lorenzo Maselli, Guy Kouarata, Jean-Pierre Donzo, Vinet Coetzee, Minique de Castro, Peter Ebbesen, Edita Priehodová, Eliška Podgorná, Viktor Černý, Susanne T. Green, Pakou Harena, Lebarama Bakrobena, Forka Leypey Mathew Fomine, Zelalem GebreMariam Tolesa, Wendawek Abebe Mengesh, Michael de Jongh, Himla Soodyall, Koen Bostoen, Chiara Barbieri, Maximilian Larena, Helena Malmström, Carina M. Schlebusch

## Abstract

Africa harbors the richest diversity of mitochondrial DNA lineages, reflecting its central role in human evolutionary history. Early studies of mtDNA variation provided the first genetic evidence for the African origin of modern humans. With complete mitochondrial genome sequencing, we can now reconstruct maternal lineages with high resolution, yet large parts of the continent remain underrepresented. Using a newly developed long-range sequencing assay, we generated 1,288 complete mitochondrial genomes from 14 countries across sub-Saharan Africa, focusing on previously understudied regions. We combined these with over 3,600 publicly available African mitogenomes to produce a comprehensive dataset and updated overview of maternal genetic diversity across the continent.

We contextualized this diversity with autosomal structure and information on major human expansions, integrating archaeological and linguistic evidence. Our analyses reveal a demographic expansion of Niger-Congo speakers around 17 thousand years ago (kya), followed by a second expansion associated with Bantuspeaking groups around 6 kya. We identify haplogroup L3e as a key marker of this early Bantu expansion, tracking its spread across sub-Saharan Africa. Distinct demographic signatures also emerge for different geographic sub-branches of Bantu speakers.

These findings highlight the power of mitochondrial DNA to trace maternal ancestry and demographic history in Africa, while also acknowledging its limitations for phylogeographic reconstruction.

## Introduction

Genetic studies of human populations have described different degrees of structure between and within continents,^1,2,3^ with patterns of relatedness often correlating with geography.^2,4,5^ This holds for both autosomal variation and for the maternally inherited mitochondrial DNA (mtDNA). MtDNA genomes are often employed for studying population origins, diversity, and migration history because of their high mutation rate, small size (16569 base pairs), uniparental inheritance, lack of recombination, and abundance in cell copies.^6^ Given its maternal inheritance, the mtDNA genome’s sequence provides valuable information on an individual’s maternal ancestry. Moreover, the relevance of mtDNA has recently been highlighted by studies of ancient DNA, as it is more easily retrievable in context of damaged and degraded DNA material.

Numerous studies have focused on one or more of the hypervariable regions of the mtDNA genome, sometimes also including haplogroup-defining SNPs. ^7,8,9,10,11^ Methods to sequence mtDNA have been available since 1977^12^ and have constantly become less complex and faster. In 1981, the complete sequence of the human mtDNA genome was published.^13^ The first full mtDNA genomes were amplified using PCRs of multiple overlapping fragments, which was recently reduced to amplification in two fragments.^14^

MtDNA sequences that are closely related are grouped together in mitochondrial haplogroups, a collection of similar mtDNA sequences that share single nucleotide polymorphisms (SNPs) inherited from a common ancestor. Mitochondrial haplogroups are conventionally named after capital letters. With multiple mtDNA genomes, it was confirmed that all modern humans carry mitochondrial haplogroups within the macro-haplogroup L, which is further subdivided into subclades L0 to L7. Notably, the highest diversity of mtDNA sequences is found in Africa, which has led to considerable scientific attention on mtDNA genomes from the region. Haplogroup L3 gave rise to all mtDNA sequences outside the African continent (M, N and R).^3,15,16^ Although mitochondrial haplogroups M1 and U6 originated outside Africa, they are generally considered African haplogroups, as they were reintroduced by back-migrations and are predominantly found in Africa.^17,18,19^

Linguistic and geographic structure often correlate with genetic structure in Africa. ^4,5,20,21^ Traditionally, four major indigenous language phyla are identified in Africa: ^22^ Khoisan, Niger-Congo, Nilo-Saharan, and Afro-Asiatic (Supplementary Figure 1). Today the genealogical validity of several of these phyla is increasingly questioned^23^ (Figure 1 from Dimmendaal, 2008^24^). However, previous genetic studies used the proposed linguistic phyla to group populations into distinct macroregions which correspond to boundaries of genetic structure, with the suggestion that speakers of related languages would be sharing a similar demographic history (Supplementary Note 1).^20,25^ In this study, we therefore use similar grouping based on anthropological, linguistic and geographic information to explore demographic history, patterns of genetic structure and signatures of expansion associated to some linguistic families.

**Figure 1:**
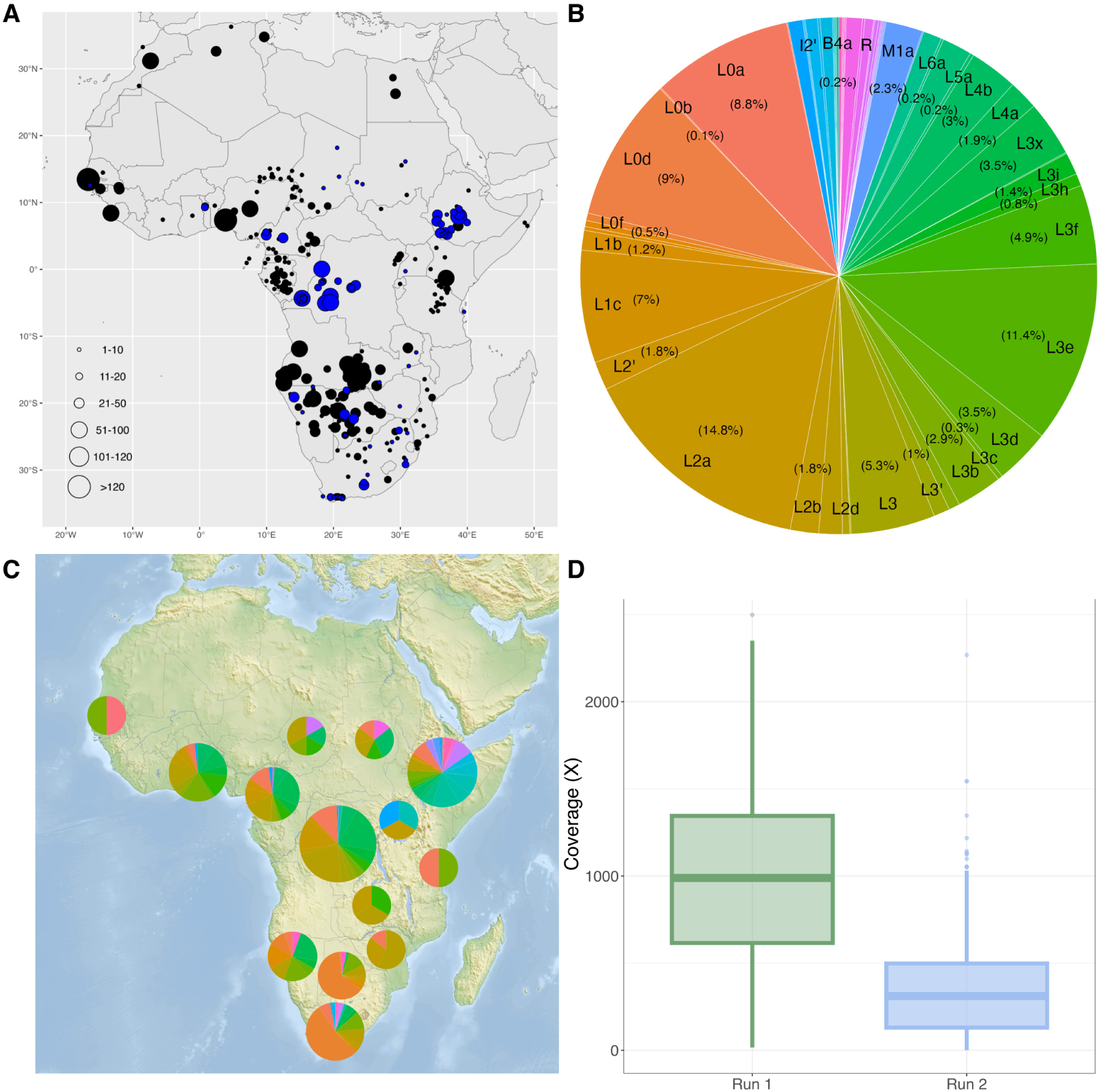
Information on sampling location, sequence coverage, and haplogroups of 1,288 newly sequenced individuals. A) Locations of the newly sequenced samples (blue) and the comparative data (black). Size of the circles corresponds to the sample size at that location. B) The frequency of haplogroups among the newly sequenced individuals. C) Pie charts showing the frequency of haplogroups among the newly sequenced individuals per country. Pie charts from countries with a higher number of individuals were depicted larger, colours correspond to those used in B. D) Boxplot of mtDNA coverage for the individuals, separated into the two sequencing runs.

One early-diverging genetic ancestry is mostly found in people speaking so-called Khoisan languages. Khoisan languages were once grouped together as non-Bantu languages sharing the extensive usage of click sounds ^22^ but comprise at least three distinct families in Southern Africa, collectively referred to as Southern African Khoisan (SAK), i.e., Kx’a (or Ju, referred to as Northern Khoisan), Tuu (Southern Khoisan), and Khoe-Kwadi (Central Khoisan), as well as two isolate languages in Eastern Africa, i.e., Hadza and Sandawe.^26^ SAK languages count roughly 250 000 speakers today. Throughout this paper, we refer to the languages as *Khoisan* and the people as *Khoe-San* (the designation preferred by the San Council).^27^ The Khoe-San comprise hunter-gatherers (San) and herders (Khoi or Khoekhoe). The Khoe-San represent one of the two branches in the earliest population divergence among *Homo sapiens*.^28,29,30,31^

Another genetic ancestry mostly represented in West-Central Africa and in most sub-Saharan Africa is associated with the Niger-Congo phylum. Niger-Congo is a phylum that includes subbranches of debated genealogical relatedness. Most classifications agree on a shared ancestry of the Volta-Congo languages, a group that includes the large Bantu family and some related languages in West-Central Africa. There are 400–600 million Niger-Congo speakers.^32,33^ Autosomal microsatellite data reconstructed an expansion associated with some Niger-Congo speakers starting around 7.4 kya,^34^ although the Niger-Congo expansion was previously associated with the stabilizing climate during the Holocene (the current geological epoch that started 11.7 kya). ^35,36^ The region of origin of the expansion of Niger-Congo speakers is currently unknown, but North Africa, as well as various regions in West Africa have been proposed.^37,38^ Newer linguistic classifications exclude Mande and Ubangian languages from the Niger-Congo phylum.^24^ Within Niger-Congo, 250 million individuals speak one or more Bantu languages.^33^ The vast spread of Bantu speakers across Africa today is due to a migration process known as the Bantu Expansion. This Bantu-speaking migration started in West Africa (Nigeria/Cameroon) around 5–3 kya.^39,40,41,42,43^

Nilo-Saharan languages are spoken in Northeast and Eastern Africa, in the upper parts of the Nile and Chari rivers.^44^ There are about 70 million Nilo-Saharan speakers today.^32^ Of the four language families proposed by Greenberg in 1963, Nilo-Saharan is among the least widely accepted.^44^ However, there are some specific genetic ancestries shared among Nilo-Saharan speakers (see Figure 5 and S19 from Tishkoff *et al*.^20^). Alongside Nilo-Saharan related ancestry in Eastern Africa, Afro-Asiatic related ancestry is another ancestry found among Eastern African populations. Afro-Asiatic languages are subdivided into six families: Berber, Chadic, Cushitic, Egyptian, Omotic and Semitic^45^ and are spoken in Northern, Northeastern and Eastern Africa, including the Horn of Africa, and in the Middle East, by 650 million people today.^32^ Apart from the indigenous language phyla that are identified in Africa, Indo-European languages like English, Afrikaans, French and Portuguese are also spoken after their introduction during colonial times.

Genetic signatures associated to specific regions and/or cultural-linguistic groups can be found also in the mitochondrial haplogroups from Africa.^1,46,47,48^ For classification of the geographic areas, consistently used throughout this study, see Supplementary Figure 2. L0d, one of the two subclades of the most deep-rooted clade of the mtDNA phylogeny,^1^ occurs primarily among the Khoe-San people in South Africa, Namibia and Botswana.^1,8,10,49,50^ Specific subgroups of L0d, namely L0d3, L0d2a and L0d1b, show higher frequencies in the south, wheareas L0d2c and L0d1a are distributed more centrally within Southern Africa.^8^ The majority of L0d subgroups shows significant signs of expansion.^8^ Individuals carrying mitochondrial haplogroup L0k generally have a more northern distribution than individuals carrying L0d lineages: L0k is also mostly found among Khoe-San individuals, in Namibia, Angola, Botswana^50^ and Zambia.^8^ L0a, on the other hand, is very common and widespread. It has been proposed that L0a originated in East Africa,^51^ but the highest frequencies of L0a are currently found in Mozambique. This distribution has been attributed to the Bantu Expansion.^51^

The expansion of Bantu speaking people is associated with various haplogroups which include L0a, ^52,53,54^ L1c,^55,56^ L2a,^46,54^ L3b,^57^ and L3e.^58,59^ Haplogroup L1 is most frequent in West and Central Africa,^60,61^ with sublineages L1b frequent in West Africa (Ghana and Ivory Coast),^62^ and L1c frequent in Central Africa (Cameroon and Gabon), especially among Western Rainforest Hunter-gatherers (RHG) (sublineages L1c1a, L1c4 and L1c5).^63,64,65^ Other sublineages of L1c (L1c1b, L1c1c and L1c2) are associated with Bantu-speaking people, showing signs of recent expansion.^63^ L2a is the most frequent mtDNA haplogroup in Africa^62^ and is prevalent in many parts of the continent, but is specifically abundant in Central Africa.^66^ Haplogroup L3 encompasses African-specific branches as well as non-African branches. L3e is the most widespread branch, and a subclade L3e1 is common among South-Eastern Bantu speakers.^67^ L3b and L3d are found mainly in West Africa and L3f is found most frequently in East Africa,^46^ where it is found in Afro-Asiatic groups. Its subgroup L3f3 is found in Chadic speakers living today in Lake Chad region.^47^ East Africa also hosts the more rare haplogroups L4– L7.^62,60^ Haplogroup L0f is characteristic in Afro-Asiatic groups. ^68^ No mitochondrial haplogroups have been specifically associated with Nilo-Saharan speaking groups, except a high proportion of L0a (otherwise associated with the Bantu Expansion), which is present among Kenyan Nilo-Saharan speakers.^68^

Despite significant advances in our understanding of human maternal history over the past decades, studies on full mtDNA genomes have frequently been limited to specific regions or populations in Africa, or have focused solely on particular mitochondrial haplogroups.^17,47,69,36,70,71,48^ When Behar *et al.*^72^ conducted a reassessment of the mtDNA phylogeny with full genomes, the geographic coverage of the African continent was limited. On the other hand, the last full overview of mtDNA diversity on the African continent performed by Salas *et al.*,^46^ has not been updated with full mtDNA genomes. Here, we present 1,288 newly sequenced full mtDNA genomes merged with 3,620 published full mtDNA genomes from the African continent, serving as a new, updated overview and summary of African maternal history. We outline the phylogenetic relationships between the 4,908 mtDNA sequences, examine their contemporary distribution, describe potential regions of origin for the most common mitochondrial haplogroups, and investigate female effective population size changes. By providing a more comprehensive and geographically inclusive analysis, this study offers novel insights into the evolutionary history and demographic patterns of maternal lineages across Africa.

## Results

We first examine the samples generated within this study and then integrate these findings into the larger continental comparative dataset. This approach allows us to compile an inclusive overview of the maternal haplogroups and ancestries across different regions and language groups in Africa, providing insights into the continent’s genetic diversity and historical migrations.

### Overview of newly generated samples

For this study, we produced complete mtDNA genomes from 1,288 new samples from understudied regions within Africa, including Central and East Africa. The individuals come from 66 sites across 14 different African countries (blue circles in Figure 1A). Full mtDNA sequences were amplified using long-range PCR and sequenced on the PacBio Sequel II platform in two sequencing runs. The average coverage for the first run, encompassing 292 samples, is 994x and the average coverage of the second run including 1,024 samples is 340x (Supplementary Figure 3). Average mtDNA coverage achieved with both PacBio Sequel II sequencing runs surpasses coverages that we achieved previously with the PacBio Sequel (87 samples with an average coverage of 292x),^73^ despite the fact that three to twelve times as many samples were pooled on the PacBio Sequel II. We filtered our data with a minimum of 30x coverage.

Haplogroups were assigned using HaploGrep3.^74^ Haplogroup composition (Figure 1B), shows that L2a (14.8%), L3e (11.4%), L0d (9.0%), L0a (8.8%), and L1c (7%) occurr most frequently. Mitochondrial haplogroups per site are shown in Supplementary Figure 5. All haplogroups were associated with a maternal ancestry, according to information available in previous studies (Supplementary Table 4) using linguistic groups or geography (Supplementary Figure 6 by site and Supplementary Figure 7 by country). We confirm high frequencies of haplogroups L0d and L0k (associated with Khoe-San ancestry) in Southern Africa (South Africa, Botswana, and Namibia); L0a, L2a, and L3e (associated with the Bantu spread) widespread across sub-Saharan Africa; L3f, L4a and L4b in East Africa (but also in appreciable frequencies in Chad and Cameroon). We note the presence of 76 sequences (6%) with non-African mitochondrial haplogroups (of Asian and European origin). The majority of these (68%) are from Ethiopia.

### Description of database containing nearly 5,000 complete mtDNA sequences

A total of 3,620 mtDNA sequences from 40 published studies were retrieved from NCBI. Mitochondrial haplogroups were assigned for all these individuals using HaploGrep3 (Supplementary Figure 8A). This was combined with the mtDNA sequences of the 1,288 newly sequenced individuals to make a total of 4,908 mtDNA sequences (Supplementary Figure 8B). From this point onward, the analyses will focus only on this combined dataset. The most common haplogroup in our African dataset is L0d (17.4%), associated with Khoe-San maternal ancestry, followed by L2a (13.7%) and L3e (12.1%), both associated with the Bantu Expansion.

To provide a general overview of the phylogeny of African mitochondrial haplogroups, we generated a Bayesian phylogenetic tree with a selection of 39 mtDNA sequences representing all major African haplogroups in the dataset up to two classification levels (e.g. L0d, L2a), with a Neanderthal mtDNA as an outgroup, and calibrated with the mutation rate of 2.285 × 10*^−^*^8^ per site per year^60^ (Figure 2). The bar charts on the right side of the figure depict the composition of broad linguistic groupings and geographic regions associated with the individuals belonging to each haplogroup tip. From the Bayesian phylogeny, we infer the matrilineal most recent common ancestor of humans and Neanderthals to date back to 305 kya [95% confidence interval (CI): 267– 344] (Supplementary Figure 9). Estimates from other studies (660^75^ and 413 kya^76^) are based on the human-chimp divergence time and a slower mutation rate, respectively. Our estimate postdates these previous estimates with 108–355 thousand years (ky). We infer the first split in the modern human mtDNA phylogeny to occur between haplogroup L0 and all other mitochondrial haplogroups around 132 kya (95% CI: 117–148). The TMRCA of individual haplogroups can be found in Supplementary Table 6.

**Figure 2:**
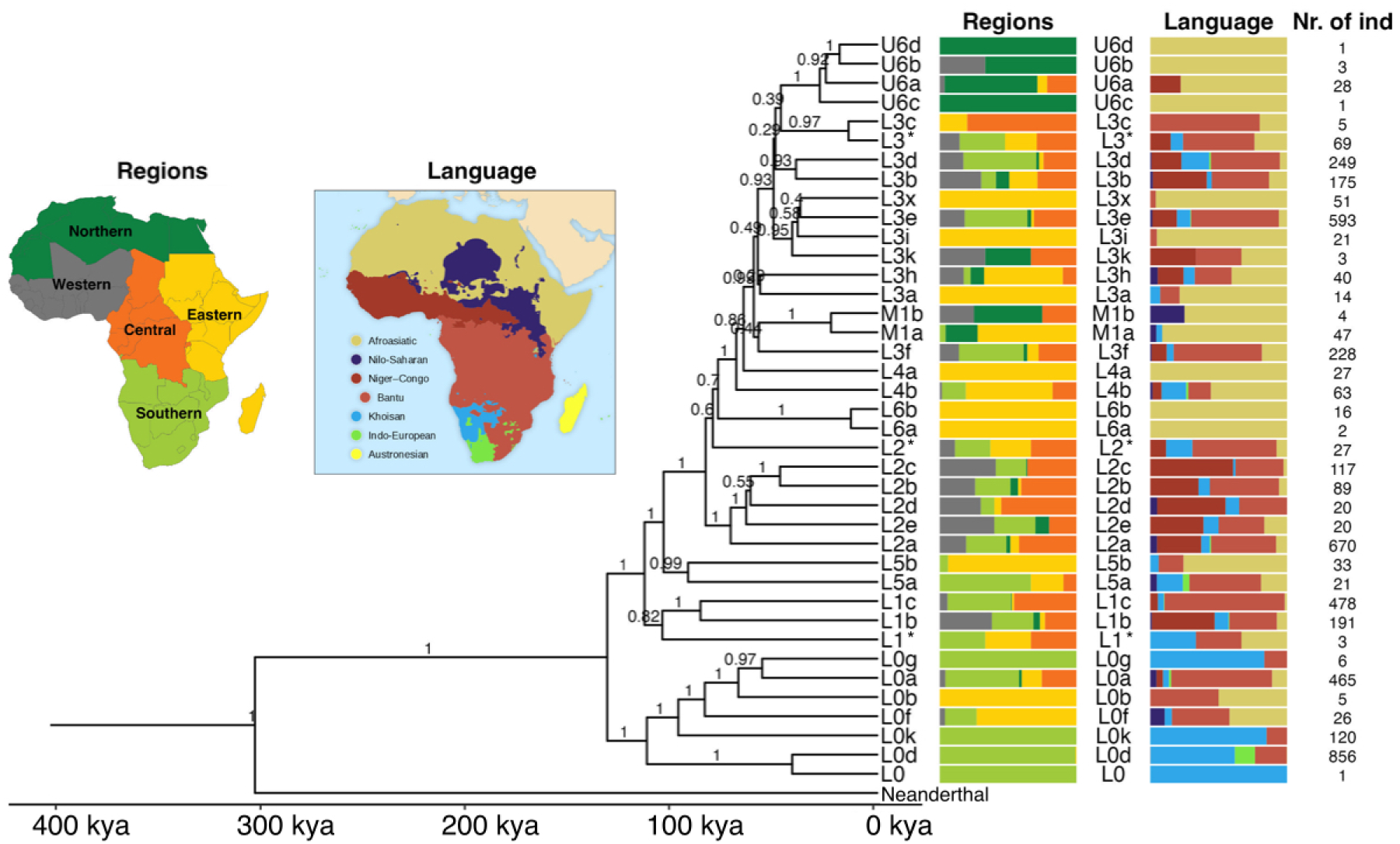
The Bayesian tree topology of African mitochondrial haplogroups based on 39 samples representing major haplogroups. On the right side of the tree, the regional and linguistic affiliation of the samples belonging to the corresponding leaves are shown. On the right, the number of samples belonging to each leaf. The inserts show the proportion of individuals according to geographic regions or language group. Posterior probability scores are shown at the nodes. Haplogroups denoted with an asterisk (*) contain all sequences not belonging to the other subhaplogroups of that clade. For example, L1* contains all L1 sequences not belonging to L1b or L1c.

In the following analyses, when referring to Niger-Congo speakers, we included only Niger-Congo speakers speaking non-Bantu languages, as well as individuals speaking Mande languages. We note that no individuals speaking Ubangian languages are present, as this linguistic branch is of disputed affiliation to the Niger-Congo phylum. These groups have a Western African to West-Central African distribution and are concentrated above the equator. Bantu-speakers have a wider distribution across sub-equatorial Africa and are analyzed as their own group in a separate analysis. In general, very few haplogroups exhibit a direct one-to-one relationship with either region or language; however, there are specific patterns that emerge and haplogroups that are present in higher frequencies in certain regions or with certain linguistic affiliations. We provide an overview of the connections between different mitochondrial haplogroups and African regions as well as language groups based on four different analyses:

1. Geographical distribution visualized on a map (Figure 3 and Supplementary Figure 10), for the ten most frequent haplogroups in our dataset, showing the current frequency distribution of haplogroups.
2. Nucleotide diversity (*π*) visualized on a map (Figure 3), quantifying the genetic variation among individuals. The nucleotide diversity of individuals carrying a specific haplogroup tends to be elevated in regions associated with the origin of that haplogroup. Thus, when applied to the mtDNA genomes of individuals sharing the same haplogroups, this analysis provides valuable insights into potential regions of origin for maternal lineages.
3. Phylogenetic relationships between the mtDNA genomes of the individuals (Figure 2 and Supplementary Figure 9 and 15–21), including information about the estimated Time to the Most Recent Common Ancestor (TMRCA).
4. Analysis of the regional distribution and languages of the individuals within haplogroups (Figure 2). Hereby, we provide an overview of which languages are spoken by individuals carrying specific mitochondrial haplogroups, as well as in which regions these haplogroups mostly occur.

**Figure 3:**
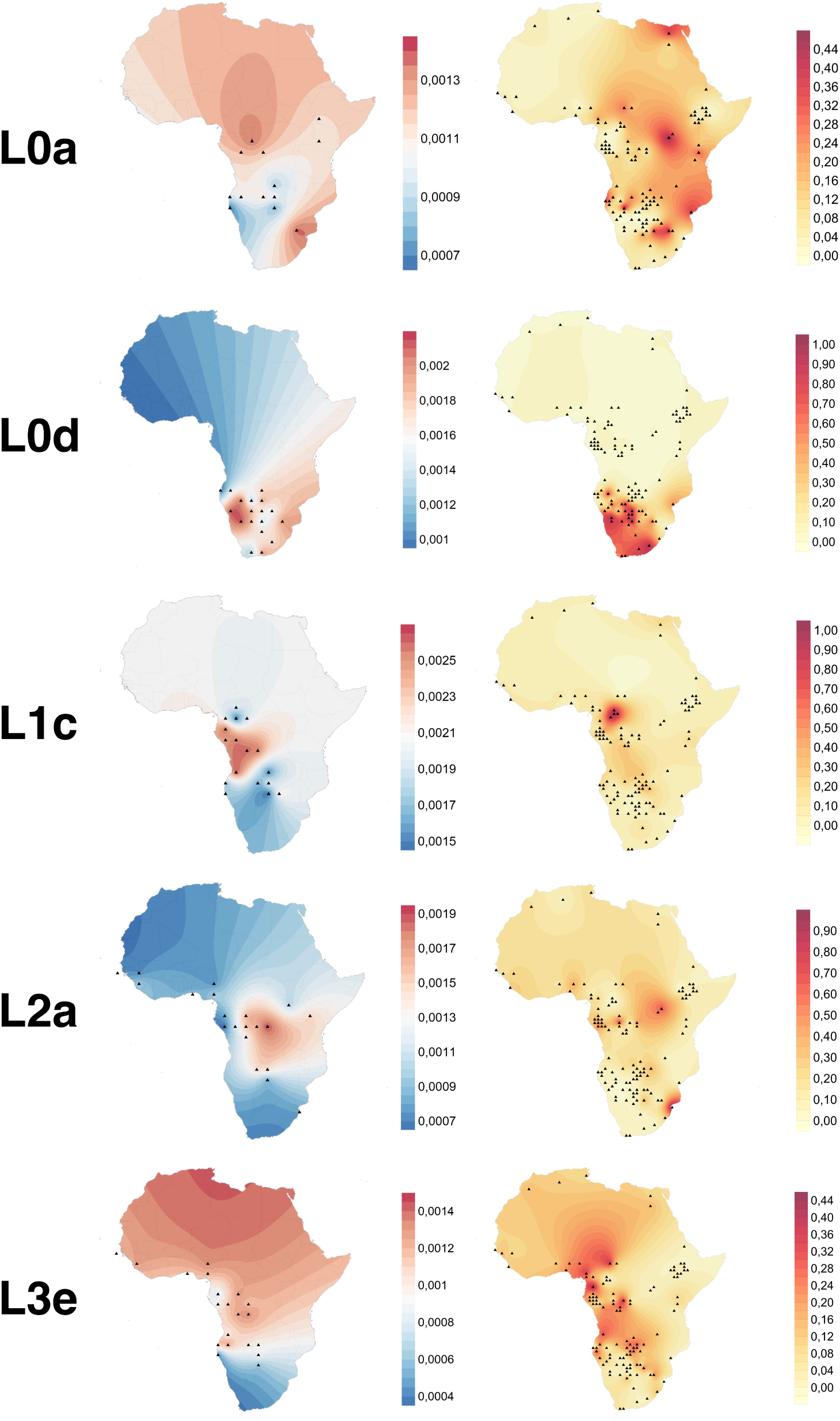
Nucleotide diversity (left) and frequency (right) maps for five most common haplogroups in our dataset (in alphabetical order: L0a, L0d, L1c, L2a and L3e) visualized on a map. Nucleotide diversity maps are based on 3 degrees bins and a minimum of 10 individuals for nucleotide diversity calculation, frequency maps are based on 1 degree bins and a minimum of 10 individuals.

#### Haplogroup descriptions

The first split in the modern human mtDNA phylogeny is between **L0** and all other mitochondrial haplogroups.

Haplogroup **L0d** is exclusively found in Southern Africa and mostly in Khoisan speakers (61.8%). A smaller proportion speaks Bantu languages (23.4%); Eastern Bantu (11.4%) and South-Western Bantu (9.9%), and the Indo-European language Afrikaans (14.7%). Most of the L0d-carrying Afrikaans speakers (*>*80%) are from Namibia,^77^ or self-identify as Coloured from South Africa. The most recent common ancestor of all L0d sequences lived 116.3 kya (95% CI: 91.9–142.8). The highest nucleotide diversity is found slightly west of the Kalahari desert (Figure 3).

**L0k**, with a TMRCA of 64.7 kya (95% CI: 47.5–82.8), has the highest occurrence in northeastern Namibia and northern Botswana, and is less widespread than L0d. The majority of the sequences belong to L0k1 (91%), and more specifically to L0k1a1 (58% of all L0k sequences). This haplogroup starts to diversify around 10–12 kya. Most of the individuals (85%) carrying haplogroup L0k speak Khoisan languages, and the remaining 15% speak Bantu languages; Eastern Bantu (10.8%) and South-Western Bantu (4.2%).

Haplogroup **L0f** is a rare haplogroup found in Eastern Africa. All L0f sequences share a common matrilineal ancestor around 82.5 kya (95% CI: 69.3–95.3) and diverges roughly 75–50 kya, earlier than L0k. Bantu speakers (34.6%) and Afro-Asiatic speakers (30.8%), make up the majority of the L0f carriers. Eastern Africa hosts mainly L0f2 (connected to Afro-Asiatic speaking individuals), whereas Southern Africa mainly hosts L0f1 (connected to Bantu speaking individuals). The single individual with haplogroup L0f from West Africa carries L0f2a.

Within L0, **L0a** (TMRCA of 67.4 kya (95% CI: 54.2–81.6) has a peculiar geographic distribution, different from the sister branches L0d, L0k and L0f. L0a is very widespread, and is found at frequencies up to 40% in the northeastern part of the Democratic Republic of the Congo (DRC), the northeastern part of South Africa, and even in northern Egypt. It is more or less absent in the very southwestern part of the African continent. Although most of the L0a carriers are Bantu speakers (73.7%), it can be found among all language groups. The highest nucleotide diversity is found in the DRC and South Africa. The remaining L0 branches **L0b** and **L0g** appear at a very low frequency, in East and Southern Africa respectively.

Following L0, haplogroup **L1** is the next haplogroup to split off from all the other haplogroups around 105 kya (95% CI: 87.8–122.5), with its subclades L1b and L1c sharing a common ancestor 86.7 kya (95% CI: 69.8–102.9). Haplogroup **L1b** is found mainly in West (38%) and Southern Africa (30.4%). L1b carriers mainly speak non-Bantu Niger-Congo and Mande languages (46.0%), and Bantu languages (34.8%), and to a small extent Khoisan languages (10.2%). Haplogroup **L1c** is among the most frequent haplogroups in our dataset, with highest frequencies in the homeland of the Bantu Expansion (Nigeria and Cameroon). It is also found in the DRC, Angola, Zambia, and Namibia. The highest nucleotide diversity is in the west coast of Central Africa. L1c carriers mainly speak Bantu languages (87.9%), and to a smaller extent Khoisan and non-Bantu Niger-Congo and Mande languages (4–6%). The TMRCA of L1c sequences is 81.7 kya (95% CI: 70.6–93.2).

Mitochondrial haplogroup **L2** splits off after haplogroup L5 (L5 discussed below) 84.1 kya (95% CI: 74.8–94.0). Interestingly, all L2 haplogroups (L2a–L2e) are characterized by linguistic and regional heterogeneity, with a broad range of linguistic affiliations and geographic coverage. Specifically, there are prominent proportions of Bantu (33.3– 50%) and non-Bantu Niger-Congo and Mande speakers (32.4–60.5%). Moreover, the highest frequencies are found in Central, Western, and Southern Africa. All the L2 haplogroups can be found in a small proportion (2–11%) of Khoisan speakers. The majority of L2 individuals in our dataset belong to haplogroup **L2a** (71.0%), with a TMRCA of 58.4 kya (95% CI: 48.5–69.1). The highest nucleotide diversity of L2a is in Central Africa. It is prevalent in many parts of the African continent, but the highest frequencies are found in northeastern DRC and Mozambique. 47.6% of people belonging to haplogroup L2a speak Bantu languages, 32.4% speak non-Bantu Niger-Congo and Mande languages.

Haplogroup **L3** has an extensive global distribution; all haplogroups outside of Africa trace back to the ancestral lineage of haplogroup L3. Thus, it plays an important role in understanding the out-of-Africa expansion. Additionally, L3 haplogroups have a wide distribution across the African continent, with presence in all five geographic regions and among all language groups. Contrary to what their names might imply, M1 and U6 are subgroups of L3. Haplogroup **L3b** is found in 3.6% of our dataset, with highest frequencies in East Africa, in Kenya specifically. Interestingly, it is found in relatively equal proportions in all five African regions, ranging from 9.7% in North Africa to 30.3% in West Africa. Both Bantu speakers and non-Bantu Niger-Congo and Mande speakers make up roughly 40% of the individuals carrying haplogroup L3b. Haplogroup **L3d** occurs at a frequency of 5.1%. 53.4% of the L3d sequences are from Southern Africa. 50.4% of L3d individuals speak Bantu languages, 22.0% speak non-Bantu Niger-Congo and Mande languages, and 20.3% Khoisan languages. Haplogroup **L3f** shows a similar distribution pattern to L3d. It is found in 4.6% of the dataset with highest frequencies in northern Namibia. Significantly, 64.5% of L3f individuals speak Bantu languages, and 45.9% live in Southern Africa. **L3e** frequency in the dataset is 12.1%. The highest frequencies are widely distributed from the homeland of the Bantu Expansion to northern Namibia. L3e is mainly found in Southern Africa (45.9%), Central Africa (31.0%), and West Africa (18.2%). The highest nucleotide diversity is in Western and Central Africa. The TMRCA of all L3e sequences is 36.5 kya (95% CI: 30.2–43.4). The majority of L3e carriers speak Bantu languages (64.2%). For haplogroup L3e1 and L3e2, we generated median joining networks (Supplementary Figure 22 and 23) to visualize the genetic relationships between haplotypes and infer evolutionary connections. For haplogroup L3e1, individuals with southern origins are located more at the edges of the network, whereas individuals with Central and Western origins are located more centrally in the network. This pattern is not observed for L3e2.

Haplogroups **L4, L5, and L6** are mostly found in Eastern Africa. Haplogroup L4a, L6a, and L6b are exclusively found in East Africa and among Afro-Asiatic speakers. L5a has a more southern distribution, with 66.7% of all our L5a sequences coming from Southern Africa. Further, 52.4% of the L5a carriers speak Bantu languages.

Although mitochondrial haplogroups **M1 and U6** originated outside Africa, they are generally considered African haplogroups, as they were reintroduced by back-migrations and are now predominantly found in Africa^17,18,19^. Haplogroups U6a, U6b, U6c, and U6d are almost exclusively found among Afro-Asiatic speakers and 66.7–100% of the people carrying these haplogroups live in North Africa. Interestingly, 33.3% of the U6b carriers live in West Africa. Haplogroup M1 shows a slightly more diverse pattern for both regions and languages when compared to U6 haplogroups. M1a, which is found in 47 samples in our dataset, occurs mostly in East Africa (72.3%) and 91.3% of M1a carriers are Afro-Asiatic speakers. These haplogroups are represented by relatively few individuals in our dataset, and more sequences would be needed to consolidate these haplogroups profiles.

### Variation in female effective population sizes associated to language groups

All individuals were assigned a language group if language information was available. We refer to Supplementary Note 1 for the use of linguistic labels for genetic clusters. Comparisons were made between Afro-Asiatic, Nilo-Saharan, Bantu, non-Bantu Niger-Congo and Mande, Khoisan, and Afrikaans (an Indo-European language of Dutch origin that developed locally in the 17th century). The distribution of mitochondrial haplogroups among these different groups (Supplementary Figure 11) illustrates the maternal lineage diversity within the African continent. Variation of female effective population sizes (*N*_e_) through time for each of these groups was determined through Bayesian Skyline Plots (BSP) (Figure 4). *N*_e_ of various Bantu speaking groups (North-Western Bantu, West-Western Bantu, South-Western Bantu and Eastern Bantu) were analyzed separately and are shown in Figure 5.

**Figure 4:**
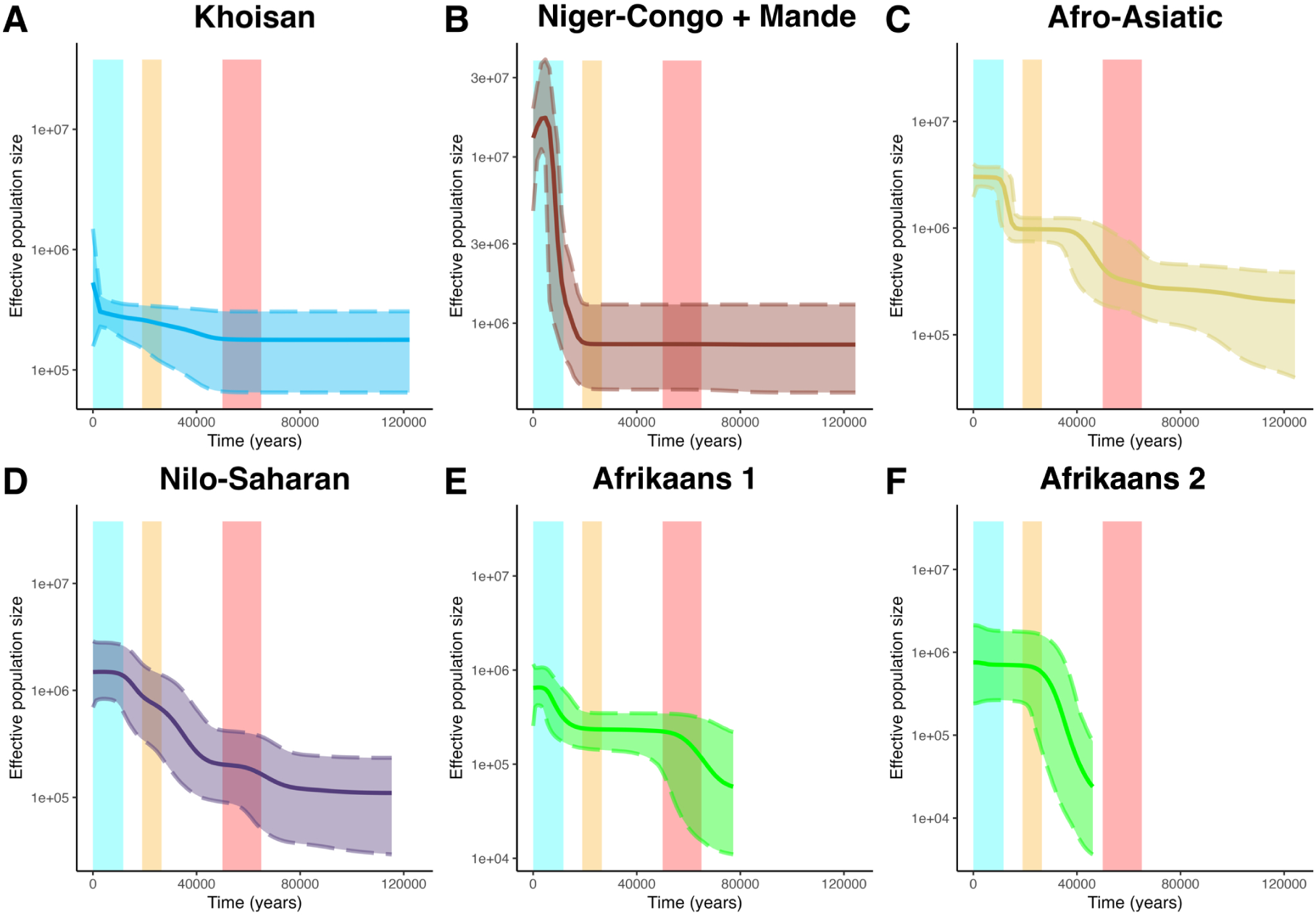
Bayesian Skyline Plots (BSP) showing variation of *N*_e_ through time for separate language groups. A) Khoisan (only L0d and L0k carriers), B) Niger-Congo and Mande (excluding Bantu speakers), C) Afro-Asiatic, D) Nilo-Saharan, E) Afrikaans (including only L0d and L0k haplogroups) and F) Afrikaans (including only haplogroups of Eurasian origin) speakers. The bold middle line represents the mean estimates and the two dashed lines represent the 95% highest posterior density (HPD) intervals. The red area denotes the time of the out-of-Africa migration (65–50 kya), the yellow area the Last Glacial Maximum (LGM)(26.5–19 kya), and the blue the Holocene (last 11.7 ky). *N*_e_ is represented with a log-scale on the Y-axis, and the years in the past are represented on the X-axis. A zoom-in of the last 25 000 years for Niger-Congo and Mande can be found in Supplementary Figure 24.

### Khoisan

Because genetic studies have been focusing intensively on Khoe-San populations due to their deep population history and distinct uniparental lineages, the dataset includes a relatively large number of Khoisan speakers (783 individuals, roughly 16.0% of the dataset). The predominant mitochondrial haplogroup among people speaking Khoisan languages is L0d (57.8%). Khoisan speakers show a slight increase in female *N*_e_ after 65–50 kya, which corresponds to the time period of the out-of-Africa expansion (Figure 4A). Figure 4A is based only on L0d and L0k haplogroups, as these were the only haplogroups found among local Stone Age hunter-gatherers before East Africans and Bantu speakers moved into the area, as shown by aDNA studies.^30,78^ For a complete analysis of the effective population size of all haplogroups currently found among Khoisan speakers see Supplementary Figure 12. We also characterized the haplogroup distribution among six different Khoisan groups (Kalahari Khoe, Khoekhoe, Kx’a, Tuu, Sandawe and Hadza)(Supplementary Figure 13), and observe different haplogroups among the Sandawe and Hadza, although only a few individuals from these groups were included in the dataset. Furthermore, it seems like Khoekhoe and Kalahari Khoe have more external input (non-L0d/k) compared to Tuu and Kx’a. It is however difficult to deduce whether this input is from Bantu-speakers or East African groups.

### Niger-Congo and Mande

Individuals speaking non-Bantu Niger-Congo and Mande languages are analyzed together as they share a similar autosomal genetic ancestry.^20^ In our analysis, we group individuals spanning a region from Senegal in the West to the Central African Republic in the East. Major haplogroups found among these individuals are L2a (28.6%), L3e (14%) and L1b (11.5%). In this group we see a population expansion starting around 17 kya, with a maximum growth around 9 kya (Figure 4B and Supplementary Figure 24). We also analyzed the female *N*_e_ of Niger-Congo and Mande speakers including Bantu speakers (Supplementary Figure 25), which shows two expansion periods, one starting around 19 kya, and the other starting around 5 kya.

### Afro-Asiatic

Afro-Asiatic speakers in our dataset mainly come from the northern part of the African continent. Although the database selection was restricted to African haplogroups, non-African haplogroups of European origin were found among the 1,288 newly sequenced mitogenomes: like R0a at 2%, K1a at 2%. The Afro-Asiatic group includes haplogroups that originated outside of Africa and returned through back-migration (M1 at 7.2% and U6 at 4.2%), as well as haplogroups associated with Western African non-Bantu Niger-Congo and Mande, and Bantu speakers (L2a (8.5%) and L0a (7.8%)). The BSP shows an increase (three- to ten-fold) after the out-of-Africa expansion, and a subsequent increase starting around 20 kya.

### Nilo-Saharan

The small number of individuals associated to the Nilo-Saharan group in our dataset (N=81) harbour a wide diversity of mitochondrial haplogroups, like L2a (39.5%), L0a (24.7%) and L3e (11.1%). Individuals associated to the Nilo-Saharan group show an increase of *N*_e_ (threeto ten-fold) after the time period of the out-of-Africa expansion, similar to Afro-Asiatic speakers but slightly more delayed. Interestingly, the BSP shows a steady increase in *N*_e_ from around 45 kya, stabilizing around 12 kya. With a relatively small sample size we do not recognize any distinct mitochondrial haplogroup associated to this group (Figure 2). This reiterates the lack of linguistic and genetic unity among the Nilo-Saharan phylum.

### Afrikaans

Afrikaans speakers in this study are only individuals self-identifying as “Baster”^77^ and “Coloured”^79^. A rather large part of the people speaking Afrikaans in Southern Africa carry the Khoe-San associated haplogroup L0d. The individuals who speak Afrikaans and carry L0d haplogroups show two *N*_e_ expansions in the BSP (Figure 4E); one preceding the time period of the out-of-Africa (65–50 kya), the other starting around 10 kya, which is similar but slightly later than that observed in Afro-Asiatic speakers. Haplogroups that originated in Eurasia, such as M and B are found in low percentages (5.4% and 2.4% respectively) in this group. The *N*_e_ changes of Afrikaans speakers carrying Eurasian haplogroups can be found in Figure 4F and shows one expansion from roughly 45 kya until the Last Glacial Maximum (26.5–19 kya). The BSP plot with all Afrikaans speakers can be found in Supplementary Figure 14 and is similar to that of Afrikaans speakers that carry L0d haplogroups (Figure 4E).

### Bantu

The largest linguistic group in our dataset consists of people speaking Bantu languages. This group comprises a large variety of mitochondrial haplogroups, with L1c (17.7%), L3e (16.6%), L0a (14.6%), and L2a (13.9%) showing the highest occurrences, followed by the Khoe-San characteristic haplogroup L0d (8.8%). The BSP for all Bantu individuals is shown in Figure 5B. An expansion (three-fold increase) starts around 5.3 kya and peaks around 1.8 kya, followed by a decrease in *N*_e_.

**Figure 5:**
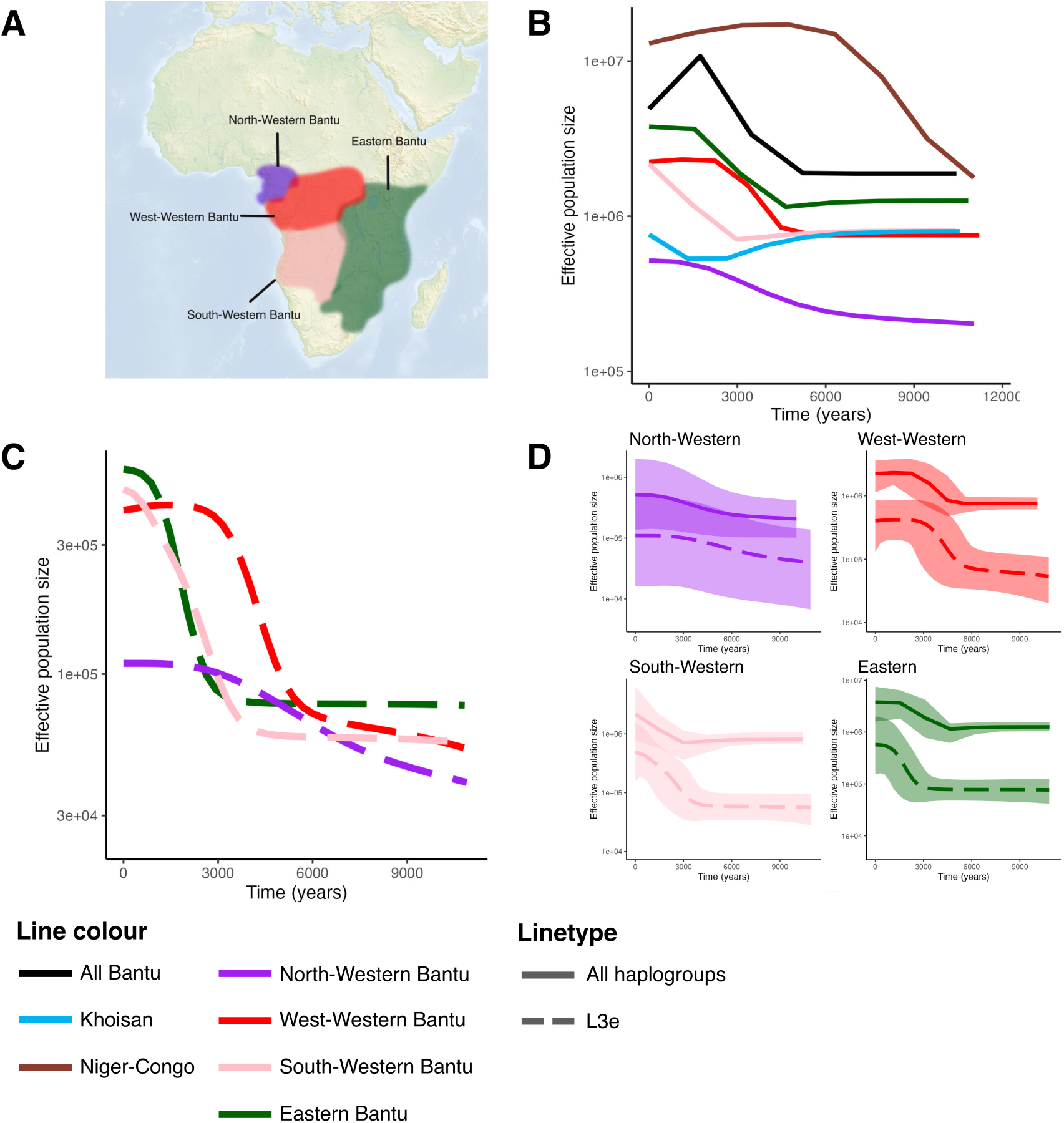
Variation of female Ne through time for separate Bantu speaking groups. A) The approximate geographic distribution of four sub branches of Bantu languages. B) *N*_e_ variation for the different Bantu speaking groups. C) *N*_e_ variation from individuals carrying haplogroup L3e in the various Bantu speaking groups. D) *N*_e_ variation from 1) all individuals within a language group (dotted line) and 2) only individuals carrying L3e within that language group. The bold middle line represents the mean estimates and the faded area surrounding the bold line represent the 95% highest posterior density (HPD) intervals. Effective population sizes are represented on a log-scale on the Y-axis, and the years in the past are represented on the X-axis.

The *N*_e_ variation was also analyzed separately for four Bantu speaking linguistic sub-groups (North-Western Bantu, West-Western Bantu, South-Western Bantu and Eastern Bantu) (Figure 5B). The patterns of *N*_e_ variation are different across these groups. Starting in and around the homeland of the Bantu Expansion, the North-Western Bantu speakers show the earliest signs of expansion: a relatively slow but steady three-fold increase, starting as early as 6 kya. The West-Western Bantu and Eastern Bantu exhibit a similar trend, both experiencing a notable rise in *N*_e_ starting around 5–4.5 kya. Both also show stabilization of this increase around 2–1.5 kya. A delayed increase in effective population size can be seen among the South-Western Bantu speakers, which starts around 3 kya and has not stabilized yet. Increases in *N*_e_ indicate that the Bantu speakers picked up more mitochondrial haplogroups while they expanded, or their *N*_e_ increased through population expansion.

Within various Bantu language groups, we analyzed *N*_e_ variation separately for the L3e individuals (Figure 5C), as the nucleotide diversity and current frequency of L3e carriers suggests that this haplogroup is associated with the earliest expansions of Bantu speakers (Figure 3). Distinct patterns can be observed over the last 6 kya, at which point they begin to converge backwards in time. In Figure 5D, we illustrate *N*_e_ variation comparing L3e against all the haplogroups of the four Bantu subgroups. Notably, for North-Western, West-Western, and South-Western Bantu groups, their population expansion comes after the one reconstructed with the L3e individuals only. However, Eastern Bantu speakers present an exception, as their expansion peak predates the one found in L3e.

The same analyses were carried for haplogroups L1c, L0a, L2a, and L3b (Supplementary Figure 26–29). *N*_e_ variation of L2a within the various language groups shows more recent expansions than all speakers of that language (Supplementary Figure 28), indicating that this haplogroup could have been picked up by expanding Bantu speaking populations. Higher nucleotide diversity values east of the Bantu homeland support this hypothesis. *N*_e_ variation of L1c carriers (Supplementary Figure 26) shows small expansions in the North-Western and West-Western Bantu speakers, but show bottlenecks in Bantu speakers living further away from the homeland (South-Western and Eastern Bantu speakers). Supplementary Figure 27 shows similar *N*_e_ variation for L0a to the rest of the individuals for each language subgroup.

## Discussion

The African continent is home to vast genetic diversity, which includes mtDNA genomes. As mtDNA is passed on through the female line, the genetic diversity in this part of the genome is informative about the female history of a population. In this study, we report a new assay-design for retrieving full mtDNA sequences using long-read sequencing. We generate novel data from understudied regions such as the DRC and Ethiopia, which is added to a newly generated database of full mtDNA sequences, allowing us to give an overview of the mtDNA diversity of the African continent. We describe different distribution patterns and reconstruct the history of the most frequently occurring mitochondrial haplogroups through frequency maps, nucleotide diversity maps, analysis of *N*_e_ variation, and time-calibrated phylogenetic trees. From the combination of these results several important findings about African maternal history came to light.

### The largest database of full mtDNA sequences to date

In a previous study, we amplified and sequenced the mtDNA genome in two fragments.^14^ Here, we present a method to sequence the full mtDNA genome with the amplification of a single fragment. It requires fewer consumables, fewer reagents, less time and greater monetary efficiency compared to all other methods to sequence the full mitochondrial genome. Additionally, we show that it is possible to sequence at least 1,024 individuals in one run, with a median coverage of 340x (Supplementary Figure 3). By sequencing 1,288 new mtDNA sequences and compiling a database containing nearly 5,000 full mtDNA sequences, this study provides the first large overview of mtDNA diversity on the African continent since 2002.^46^ We have used published data to associate mitochondrial haplogroups with maternal ancestries described by geography, culture and language. The 1,288 new mtDNA sequences, which include underrepresented regions of Central and East Africa, confirm the generic haplogroup structure and the affiliations with linguistic groups or geographical regions proposed (Supplementary Figure 7).

In this study, we generated data from 14 different countries across sub-Saharan Africa, with a significant number of samples originating from the Democratic Republic of the Congo (DRC, N=559) and Ethiopia (N=356). These regions have been historically understudied, and our research sheds light, for the first time, on the mtDNA diversity of the populations in these countries. In the DRC, individuals predominantly carry haplogroups commonly associated with Bantu-speaking populations and West-Central Africa, including L0a (11.6%), L1c (13.8%), L2a (21.8%), and L3e (19.7%). In contrast, in Ethiopia the East African haplogroups are more frequent, such as L3x (12.6%), L4a (6.7%), L4b (8.1%), L5b (6.7%), and M1a (8.1%). L3x is an otherwise rare haplogroup; the highest frequency was previously detected in Ethiopia at 6% in a Cushitic group. ^80^

It should be noted that the frequencies of the various haplogroups in our dataset are influenced by sampling bias. As an example, the most frequent haplogroup in the compiled dataset is L0d (17.4%). However, L0d is common among Khoe-San people, which are known to have a census size much smaller than Bantu speakers.^81^ Thus, the high percentage of L0d mtDNA sequences in our dataset is most likely a result of the interest in the Khoe-San people because of their unique placement in the phylogeny of all modern humans. In fact, 18.6% of the individuals in the dataset speak Khoisan languages.

### Haplogroup distribution

The majority of Khoisan speakers carry haplogroup L0d and L0k. Haplogroup L0d reaches a maximum frequency of 100% at some sites, whereas L0k frequency is much lower, maximum 33%. The distribution of L0k is also more geographically confined than L0d, which has previously been shown by Barbieri *et al.*^50^ However, Barbieri *et al.* observed peaks of L0k in northern Zambia, which we do not detect, probably due to a more extensive dataset used here. Afro-Asiatic speakers have been associated with haplogroups L0f and L3f. ^68,47^ We show that these haplogroups do occur among Afro-Asiatic speakers in our dataset, but in low frequencies (1.3% and 6.4% respectively). Haplogroup L3f additionally shows high frequencies among Bantu-speaking populations in northern Namibia. The haplogroups L0a and L2a, which are common among Bantu speaking populations, also occur at lower frequencies among Afro-Asiatic speakers. Nilo-Saharan speakers, on the other hand, show higher frequencies of some of the haplogroups also associated with Bantu speakers; L2a, L0a and L3e (totaling to 75.3%). L0a and L3e have broad distributions across the African continent, whereas L2a distribution is slightly more restricted. Mande speakers and Niger-Congo speakers whose languages are not Bantu show relatively high proportions of L2a (28.6%), L3e (14%) and L1b (11.5%).

Individuals self-identifying as Baster and Coloured speaking Afrikaans (an Indo-European language) from South Africa are characterized by a high proportion of the Khoe-San haplogroup L0d (75%). This emphasizes the impact of historical events such as colonial movements, migration, and intermarriage on the complex genetic landscape of the region.^73,79^

We also note the presence of non-African mitochondrial haplogroups (Asian and European) in various African countries, which can be attributed to either back-migration in the cases of Ethiopia and Sudan, or recent colonial influence in the cases of South Africa, Namibia and Botswana.

### The onset of the Bantu Expansion: timing and mitochondrial haplogroups

The expansion of Bantu speaking people was one of the most significant population movements in African history and had profound impacts on the demographic, linguistic, and cultural landscape of the continent.^5^ Studies based on linguistic evidence have proposed an expansion onset around 5 kya, or slightly earlier.^82,83^ The start of this expansion has been dated with autosomal microsatellite data to 5.6 kya.^34^ Further studies investigated population expansions within distinct Bantu speaking groups using admixture dating methods and IBDNe, a method to estimate *N*_e_ from the number of shared ancestors with identity-by-descent (IBD) segments.^5,84,85,86^ However, IBDNe does not reliably reconstruct relationships over 50 generations (∼1.5 ky).^87^ We have reinvestigated the timing and magnitude of the expansion using a Bayesian approach on mtDNA genomes. The expansion starts earlier for Western Bantu speakers (6 kya), with a slow but steady three-fold increase and starts later in Eastern and Southern Bantu (4.5–3 kya). This early expansion predates the estimates from microsatellites by 400 years, and those from linguistic evidence by 1000 years. However, it is important to note that this estimate of 6 kya heavily relies on the mutation rate used (2.285×10*^−^*^8^ per site per year),^60^ which is based on an average of mtDNA mutation rates surveyed in the literature. The ranges found in the literature span from 1.665 × 10*^−^*^8^ ^88^ to 2.67 × 10*^−^*^8^ ^89^. Consequently, when calculating this through to a range in year estimates, it comes down to roughly 8.25–5.15 kya. We observe a decrease in *N*_e_ of all Bantu individuals from 1.8 kya (black line in Figure 5B). This population decline should be regarded with caution, as it is most likely an artefact of population structure.^90^ In fact, this decline is not observed with the BSP analysis for different Bantu subgroups, suggesting that recent population substructure becomes apparent when merging the subgroups in a large metapopulation of Bantu speakers.

It is also interesting to investigate the specific genetic signals associated with the Bantu Expansion. The identification of haplogroups linked to the initial dispersal of Bantu speaking peoples remains a subject of ongoing investigation. We propose that L3e was one of the haplogroups associated with the earliest expansions of Bantu speakers. The nucleotide diversity observed among L3e carriers shows a north-south cline, with higher nucleotide diversity in West-Central Africa (Figure 3). Current L3e distributions are high in West-Central Africa, and in regions where Western Bantu speakers live. Within the Bantu speaking individuals carrying haplogroup L3e, separate analyses of *N*_e_ variation were conducted for each of the four subbranches of the Bantu language tree (Figure 5C and D). The BSPs of L3e lineages exhibit distinct patterns over the last 6 kya, at which point they begin to converge backwards in time. This convergence suggests a scenario wherein these L3e lineages might have been contained within the initial Bantu source population until 6 kya, potentially among the people residing in the Bantu Expansion homeland. As a caveat, it is important to be careful with the interpretation of *N*_e_ variation through time with regards to specific mitochondrial haplogroups. Mitochondrial haplogroups do not represent populations and could have entered a population at any time-point in the past. The BSP of a haplogroup before entering the studied population therefore provides no information on the female *N*_e_ of that population. Moreover, the more individuals carrying a specific haplogroup within a language-group, the more the BSP of that language-group will resemble that of the haplogroup.

We also generated median joining networks for haplogroup L3e1 and L3e2 to visualize the genetic relationships between haplotypes and infer evolutionary connections (Supplementary Figure 22 and 23). For haplogroup L3e1, individuals with Southern African origins appear more at the edges of the network, especially within subhaplogroups L3e1d, L3e1e, and L3e1a2. In contrast, individuals with Central African origins, such as those from the DRC and Cameroon, are more commonly found at the network edges in subhaplogroups L3e1a1 and L3e1a3a. This pattern suggests that these subhaplogroups likely reflect signals of back-migrations, but only for specific clades. For example, extensive back migrations of Bantu-speakers from the South to the North have occurred during the Mfekane migrations associated with unrest and displacement during the rise of the Zulu civilization. ^91^

The variation in *N*_e_ was also investigated for separate haplogroups associated to Bantu speakers (L0a, L1c, L2a, L3b), within the four Bantu subgroups (Supplementary Figure 26-29). L2a shows more recent expansions in the separate Bantu subgroups, than in the Bantu metapopulation (Supplementary Figure 28), indicating that this haplogroup could have been picked up by expanding Bantu speaking populations after the start of the Bantu migration, that the lineage itself only started to expand at a later time within the Bantu population, or that it was part of a secondary expansion wave. The onset of the *N*_e_ expansion in L0a, L1c and L3b carriers occurs at the same time in the four Bantu subgroups and in the general Bantu metapopulation. Expansion signals associated to haplogroups L2a and L3e appear later among Eastern Bantu speakers compared to the overall population of individuals speaking Eastern Bantu languages.

### Geographical mismatches between patterns of frequency and nucleotide diversity hint at high mobility of maternal lineages

By comparing the distribution of haplogroup frequencies and nucleotide diversity, we tentatively search for signatures of shifts in spatial distribution (Figure 3). For haplogroup L0a, the regions with highest levels of nucleotide diversity (western DRC and the East African coast) do not correspond to the regions with highest frequencies (eastern DRC, Mozambique, North-East South Africa, and Egypt), suggesting a shift eastwards. Even though L0a was previously proposed to have an Eastern African origin^51^, the nucleotide diversity and frequency distribution (Figure 3) here point towards an Central African origin, which is more in line with a Bantu origin - this haplogroup is also found at high frequency among Bantu-speakers. For L0d, there is high spatial overlap between the highest nucleotide diversity and the highest frequency. This could be explained by the long geographic continuity and isolation of Southern African Khoe-San groups.

L1c (Figure 3) is frequent in the homeland of the Bantu Expansion^64^ but the nucleotide diversity is highest toward the south (Angola and southern DRC). Inversely, where L1c shows high frequencies the nucleotide diversity is lowest. We hypothesize that this haplogroup, of which subhaplogroups are present in high frequencies in RHG populations, was more widespread in the past among RHG, and that nucleotide diversity has been reduced through drift and isolation there. Haplogroup L2a has highest nucleotide diversity in the center of the DRC, but highest frequencies in eastern DRC and Mozambique. It is possible that the L2a haplogroup was picked up by the Bantu Expansion and spread eastwards with subsequent local expansions.

We acknowledge the limitations of utilizing nucleotide diversity calculated in present-day human populations as a metric for tracing the origin of a maternal lineage. In instances of complete population replacement in the region of origin, no orignal nucleotide diversity information can be found, which may result in the identification of other regions as the potential origin for that haplogroup. Moreover, the extrapolation of nucleotide diversity in regions without individuals with that haplogroup using the kriging method has to be interpreted with caution.

### The onset of the expansion of Niger-Congo and Mande speakers predates previous estimates

The expansion of haplogroups associated to non-Bantu Niger-Congo and Mande languages (Figure 4 and Supplementary Figure 24) starts around 17 kya and experiences maximum growth around 9 kya, predating previous estimations based on microsatellite data (7.4 kya)^34^ and a subset of mtDNA genomes from populations in Burkina Faso (10–12kya).^36^ Although the expansion of Niger-Congo speakers was previously thought to be connected to the stabilizing climate during the Holocene (the current geological epoch that started 11.7 kya)^35^ this signal is not replicated with our data. Instead, we see a stronger impact of the end of the Last Glacial Maximum (LGM) (26.5–19 kya), in driving a demographic expansion. This signal of expansion is very deep in time and it is difficult to associate it with the origin of the language family. Linguistic reconstructions beyond 10 kya are generally not easy to infer due to the rapid rate of language evolution. This deep expansion connected to a similar genetic ancestry for the region^20^ could provide a common demographic substrate for linguistic subbranches (e.g. Mande) which have more “weak” genealogical connections to the rest of Niger-Congo family.

## Conclusion

This study provides the most comprehensive overview to date of mitochondrial DNA diversity across sub-Saharan Africa, combining 1,288 newly sequenced mitogenomes with over 3,600 publicly available sequences. By targeting understudied regions and integrating linguistic, geographic, and archaeological data, we uncover complex patterns of maternal ancestry and demographic history, including refined timings for the expansions of Niger-Congo and Bantu-speaking populations. Haplogroup L3e emerges as a key maternal marker of early Bantu dispersals, and regional differences in demographic trajectories among Bantu subgroups reveal the layered nature of population movement across the continent. Importantly, the extensive and geographically inclusive mitogenome dataset generated here serves as a valuable reference for future genetic, archaeological, and anthropological research. It offers a framework for comparative analyses, facilitates the interpretation of ancient DNA, and enhances our understanding of human population structure and migration in Africa. While mitochondrial DNA has limitations, its strength in capturing maternal lineages makes it a powerful tool, especially when supported by a reference database of this scale and resolution. This reference dataset will aid future efforts to infer African demographic history, inform ancient DNA interpretations, and improve the resolution of maternal lineage studies on a continental scale.

## Acknowledgements

We are grateful to all the individuals who voluntarily participated in this research. The authors would like to acknowledge support of the National Genomics Infrastructure (NGI)/Uppsala Genome Center and UPPMAX for providing assistance in massive parallel sequencing and computational infrastructure. Work performed at NGI/Uppsala Genome Center has been funded by RFI/VR and Science for Life Laboratory, Sweden. The computation and data handling were enabled by resources provided by the Swedish National Infrastructure for Computing (SNIC) at Uppmax partially funded by the Swedish Research Council through grant agreement no. 2018-05973. This project was supported by funding to CS from the European Research Council (ERC) under the European Union’s Horizon 2020 research and innovation programme (grant agreement No. 759933), the Knut and Alice Wallenberg foundation, the Leakey foundation and the Erik Philip Sorensson foundation. VC^̌^ was funded by Czech Academy of Sciences award Praemium Academiae. CB was supported by the URPP “Evolution in Action” of the University of Zurich and by the NCCR Evolving Language, Swiss National Science Foundation Agreement #51NF40 180888.

## Declaration of interests

The authors declare no competing interests.

## Data and code availability

All data generated or analyzed during this study is included in this published article, its supplementary information files and publicly available repositories. The generated mitochondrial data will be available for academic research use through the NCBI database with accession numbers xxx–xxx. Scripts are available on Github (https://github.com/imkelankheet/Full mitochondrial genomes).

## Declaration of generative AI and AI-assisted technologies in the writing process

During the preparation of this work the authors used ChatGPT 4o in order to assist with language and grammar checks. After using this tool, the authors reviewed and edited the content as needed and take full responsibility for the content of the publication.

## Supplementary materials

### Supplementary Note 1: The use of linguistic labels for genetic clusters

In this study, we utilize linguistic labels to refer to genetic clusters among populations. Joseph Greenberg originally proposed four linguistic families (or phyla) in 1963:^22^ Afro-Asiatic, Nilo-Saharan, Niger-Congo, and Khoisan. However, the genealogical validity of some of these phyla has been questioned by several linguists.^24,23^ Among them, Khoisan and Nilo-Saharan are the least accepted. Khoisan languages, initially considered a single family, are currently recognized as three distinct families (Kx’a, Tuu, and Khoe-Kwadi) and two language isolates (Sandawe and Hadza). The use of click consonants and linguistic borrowing led Greenberg to group these languages together as one linguistic family. For the traditional Nilo-Saharan family, constituent families are recognized but the overarching classification as Nilo-Saharan remains contentious. The unity of Niger-Congo languages is also debated, with Mande and Ubangian languages now excluded from this group. The core Niger-Congo families, including Bantu, Bantoid besides Bantu, West-Benue-Congo, Kwa, Kru, Senufu, and Gur, however remain assigned to this phylum.^38^ Afro-Asiatic is more widely accepted as a language family, and includes Berber, Chadic, Cushitic, Egyptian, and Semitic languages.

Despite the linguistic obsolescence of the labels Nilo-Saharan, Niger-Congo, and Khoisan, genome-wide genetic analyses by Tishkoff *et al.*,^20^ have demonstrated that populations grouped under these four linguistic labels proposed by Greenberg form distinct genetic clusters. Furthermore, the last full overview of mtDNA diversity on the African continent was published by Salas *et al.* and uses the four language phyla proposed by Greenberg.^46^ With a need to give an understandable label to the observed genetic clusters, and for continuity with the Salas *et al.* paper, we employ these historical linguistic labels as practical identifiers for genetic studies, acknowledging their genetic validity while recognizing their debated or outdated linguistic basis. This approach allows us to leverage historical classifications that in themselves were based on linguistic, anthropological and geographical deduction to explore meaningful genetic distinctions among populations.

A likely reason for the link between Greenberg’s classifications and genetic research is that Greenberg’s classification is primarily based on language areas rather than strict genealogical relationships. As a result, Greenberg’s approach may give the misleading impression that there is a direct correlation between genes and languages, when in reality, the correlation is more closely tied to geography. This geographic correlation explains why Greenberg’s classification might show a stronger association with genetic patterns than more recent linguistic classifications, which focus on the genealogical lineage of languages. It is important to recognize that the discrepancies between genetic data and these newer linguistic classifications do not imply that these classifications are less valid than Greenberg’s. As we choose to work with Greenberg’s classification to label genetics clusters, it is crucial to explicitly acknowledge that it highlights geographic correlations rather than linguistic genealogical ones, and that we merely use the historical linguistic labels as practical identifiers for genetic studies, due to the lack of purely genetic labels.

## Methods

### Sampling and long-read sequencing

Saliva samples or already extracted DNA samples were obtained from 1308 African individuals from 66 different sites spread over 14 different African countries (Botswana, Cameroon, Chad, DRC, Ethiopia, Namibia, Senegal, South Africa, Sudan, Togo, Uganda, Zambia, Zanzibar, and Zimbabwe). Participants donated saliva samples with written informed consent. Biological samples for this study were in part supplied by our collaborators who obtained the original ethical permission for the sampling in African countries. Supplementary Table 1 contains the ethics reference numbers and local permission details associated with these samples. Saliva samples were obtained using an Oragene DNA OG-500 kit. DNA was extracted using the prepIT L2P extraction protocol.

Primers were designed to amplify the full mtDNA genome using the program Snapgene Viewer (version 5.0.8). Eight sets of primers were tested for their ability to amplify the full mtDNA genome. The best set is shown in Supplementary Table 2. Barcoded primers were used and the full primer sequences can be found in Supplementary Table 3. A total of 1,024 unique combinations of barcoded forward and reverse primers can be created using these 32 forward and 32 reverse primers, allowing the pooling of 1,024 samples at the same time for one sequencing run. Using a unique barcode combination for every sample, we performed a PCR to amplify the whole mtDNA genome (30x (98 °C, 10 sec; 67 °C, 15 min); 4 °C ∞), (300 ng DNA, 2.4 nM primers, 200 µM of each dNTP, 1x PCR buffer and 1.25 U Takara GXL Taq in a total volume of 25 µl). Speci-ficity of PCR products was confirmed on a 1% agarose gel and purified with AMPure XP beads (Beckman Coulter). Concentrations of the cleaned PCR products were measured (Qubit, Broad Range kit) and samples were pooled (100 ng/sample). Pools were purified with 0.5x volumes AMPure XP beads and eluted in 10 mM Tris-HCl, pH 8.5. Concentration of the cleaned pools was measured on the Qubit. The complete mtDNA genomes were sequenced in two separate sequencing runs using long-read sequencing technology on the PacBio Sequel II instrument at the National Genomics Infrastructure, SciLifeLab in Uppsala. The first batch contained 292 samples, and the second batch contained 1,024 samples. From the 1,316 newly sequenced samples, there were 16 technical duplicates and 12 samples that did not yield sufficient mtDNA reads, leaving 1,288 samples for analysis.

### Haplogroup assignment

Demultiplexing of the sequencing data was performed by Uppsala Genome Centre (UGC) at NGI-SciLifeLab using the SMRT analysis pipeline. As the reads span the beginning of the Revised Cambridge Reference Sequence (rCRS), the full mtDNA sequence reads were mapped to a duplicated rCRS to create BAM files. Two bioinformatics methods to call variants were compared; DeepVariant^92^ in combination with bcftools consensus (version 1.12) and GATK HaplotypeCaller.^93^ All scripts are available at https://github.com/imkelankheet/Full mitochondrial genomes.git. The two methods were compared using vcftools (–gzdiff –diff-site). Differences among the called nucleotides between these two methods were investigated and DeepVariant was chosen for downstream analyses due to its superior reliability in detecting insertions and deletions. Average sequencing coverage was determined per sample using samtools depth (samtools version 1.12). Mitochondrial haplogroups were assigned using HaploGrep3.^74^ To ensure the independence between HaploGrep3 quality scores and coverage, we conducted a comparison by plotting the HaploGrep3 quality score against the coverages (Supplementary Figure 4).

### Database of full mtDNA sequences

A comparative database with 3,620 publicly available full mtDNA sequences was assembled. Only haplogroups associated with African populations were considered (L, M1 and U6). Sampling locations were retrieved from the publications or through correspondence with the author. In cases where precise sampling locations were not explicitly provided, but the population was known, an approximate location was deduced. This estimation involved identifying the midpoint of the known inhabited area from public databases such as the linguistic collection of Glottolog, serving as a practical proxy for the sampling location. Sequences were acquired using batch download from NCBI. The Human Genome Diversity Project(HGDP), Simons Genome Diversity Project (SGDP) and 1000 genomes project (KGP) fasta sequences were acquired from CRAM files using *bcftools mpileup*, *bcftools call* and *vcfutils.pl*. Mitochondrial haplogroups were assigned ex novo using HaploGrep3^74^ and samples were assigned a maternal ancestry based on their haplogroup, using previously published literature on haplogroup-ancestry associations (see Supplementary Table 4). All individuals were assigned to language group if language information was available. A distinction was made between Afro-Asiatic, Nilo-Saharan, Bantu, non-Bantu Niger-Congo and Mande, Khoisan and Afrikaans (an Indo-European language of original Dutch origin).

### Haplogroup frequency maps

Haplogroup frequencies were computed based on the complete mtDNA sequence database and plotted on a map of Africa using the Kriging method.^94^ As sites with a low number of individuals will bias our frequency distribution maps, we combined sites that were within one degree latitude and longitude in distance. Thereby the number of unique sites was reduced from 371 to 223. This was done to increase the sample size and thereby gain more accuracy about the haplogroup composition. Merged sites with fewer than 10 individuals were removed from the analysis; this resulted in 114 different sites. Mitochondrial haplogroup frequencies were calculated for these 114 different sites and their distribution was plotted on a map of Africa using the Kriging method using Globemapper from Blue Marble Geographics (version 22.0) and Surfer from Golden Software (version 12.8.1009).

### Nucleotide diversity maps

We computed nucleotide diversity (*π*) within the complete mtDNA sequence database, with a requirement of a minimum of 10 individuals for calculation. To ensure as many coordinates to reach this minimum number of individuals of a certain haplogroup per coordinate, we created binned sites within three degrees in latitude and longitude. This broader binning in comparison to the haplogroup frequency analysis was necessary because nucleotide diversity calculations are restricted to individuals carrying a specific haplogroup, whereas haplogroup frequencies are calculated on all haplogroups together. Therefore, a larger bin size for the nucleotide diversity helps to ensure that we do not lose too many sites due to insufficient sample sizes. Nucleotide diversity was then determined for haplogroups L0a, L0d, L1c, L2a, and L3e at each of these binned coordinates. Nucleotide diversity was visualized on a map of Africa using Globemapper from Blue Marble Geographics (version 22.0) and Surfer from Golden Software (version 12.8.1009) using the Kriging method.

### Female effective population sizes through time

For visual representation of *N*_e_ over time, Bayesian Skyline Plots (BSPs) were generated for various language groups: Nilo-Saharan, Afrikaans, Afro-Asiatic, Khoisan (all haplogroups), Khoisan (only L0k and L0d), Niger-Congo (Bantu speaking individuals not included) and Mande, Niger-Congo and Mande (300 Bantu speakers included), Bantu, North-Western Bantu, Eastern Bantu, West-Western Bantu and South-Western Bantu, as well as for various groups of individuals carrying Bantu-haplogroups among the North-Western Bantu, Eastern Bantu, West-Western Bantu and South-Western Bantu. One hundred twenty-five individuals could not be assigned a language group due to the lack of information, and they were not used for the estimation of female *N*_e_. Input files were prepared with BEAUTi using fasta files and subsequently processed with BEAST (version 1.8.4)^95^ using the Markov Chain Monte Carlo (MCMC) sampling algorithm. Details regarding the number of MCMC iterations, the burn-in iterations discarded, and the number of replicates for each language group are provided in Supplementary Table 5. A general time-reversible (GTR) substitution model was used, as suggested by ModelTest-NG version 1.0.0. A strict clock model was applied with mutation rate (*µ*) of 2.285×10*^−^*^8^ per site per year as used in Maier *et al.*,^60^ which is based on an average of mtDNA mutation rates surveyed in the literature.^96,88,89,97,98,99,100^. These estimates are based on the human-chimp divergence time, aDNA data and phylogeographic differences. A Coalescent Bayesian Skyline model was chosen, with four groups and UPGMA starting tree. The number of iterations to be discarded as burn-in was determined by examining when the likelihood had stabilized. Resulting log and trees files were merged using the program LogCombiner (version 2.6.7) and all ESS values were checked to be above 200. Visualization of the BSP was done in Tracer (version 1.7.2)^101^ and R (version 1.2.5033).

### Phylogenetic trees

Bayesian phylogenetic trees (Maximum Clade Credibility trees) were constructed using mtDNA sequences associated with African haplogroups. The Neanderthal mtDNA genome (GenBank accession number: AM948965) was included as an outgroup. One representative sample with the highest HaploGrep3 score was selected for each African haplogroup at the “L0d” level. The tree construction was performed using BEAST (version 1.8.4).^95^ The BEAST settings mirrored those described for the BSPs, employing a birth-death model. Five replicates of 100 million iterations were executed, with a burn-in of 10 million iterations. The resulting log and tree files were merged using LogCombiner (version 2.6.7). Subsequently, all effective sample size (ESS) values were verified to be above 200. Tree annotation was accomplished using TreeAnnotator from the BEAST package, and the final tree was visualized using the ggtree package ^102^ in R. For each haplogroup leaf, data on region and linguistic information were computed and visualized using R. Moreover, we generated Maximum Clade Credibility trees using BEAST for L0a, L0d, L0f, L0k, L1c, L2a, and L3e. Trees were annotated using TreeAnnotator (version 2.6.7) and these consensus trees were visualized in FigTree (version 1.4.4). L0d was added as an outgroup for all haplogroups, except for L0d itself, for which L0k was chosen as an outgroup.

### Phylogenetic networks

Median Joining Networks were constructed using Network v.10.2 (www.fluxus-engineering.com). Maximum parsimony post-processing was performed using the Steiner maximum parsimony algorithm. The epsilon parameter was left unchanged and transversions were weighted three times the weight of transitions. Networks were visualized in Network Publisher and individuals were coloured based on their regional affiliation.

## Supplementary Figures

**Supplementary Figure 1:**
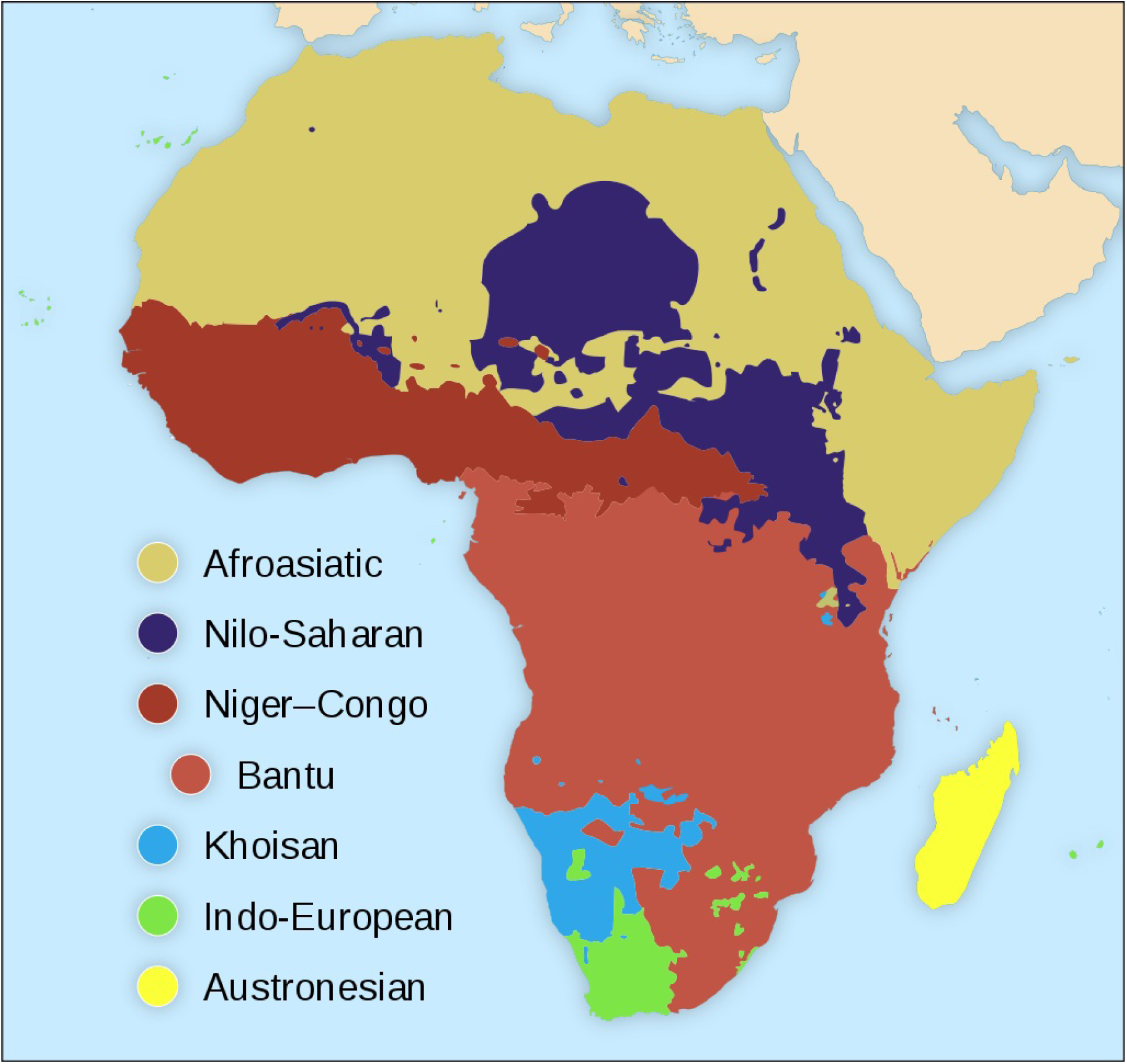
Geographical distribution of major language phyla spoken in Africa (taken from Languages of Africa, licensed under CC BY-SA 4.0.). Figure corresponds to Figure 4.1 of The Oxford Handbook of African Archaeology^103^ and was proposed by Greenberg in 1963.^22^ The language group assignment in this study is not based on the geographical information shown here; rather, it was done with careful assessment of languages spoken by the individuals, based on associated literature. These figures are meant to be used as a general overview of the distribution of language groups and language families. The language classification “Austronesian” as provided in A is not used throughout this paper.

**Supplementary Figure 2:**
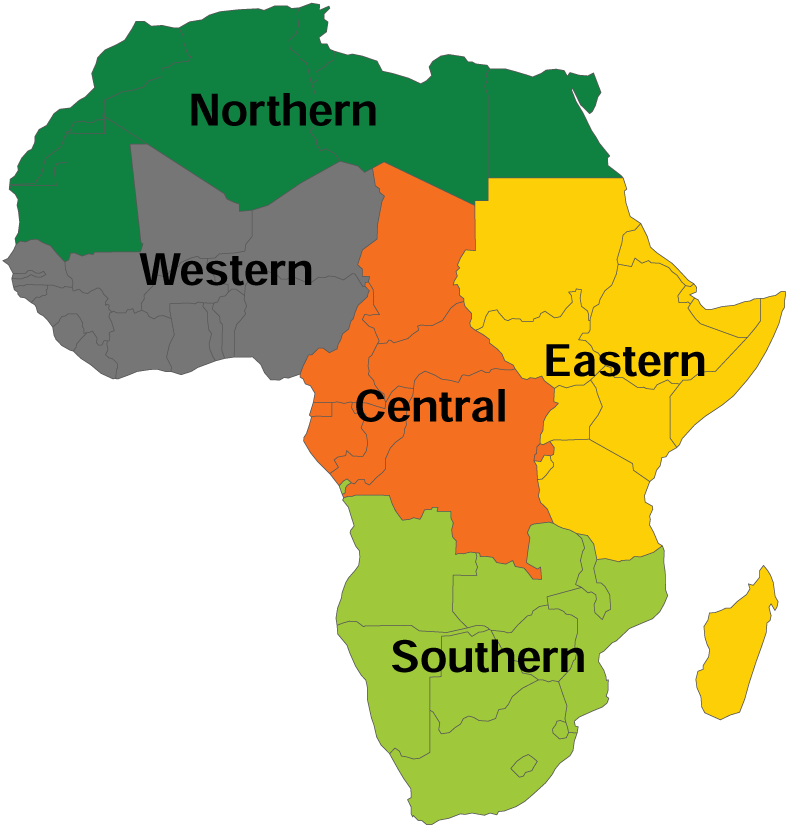
Classification of geographical areas into Northern, Western, Central, Eastern and Southern Africa.

**Supplementary Figure 3:**
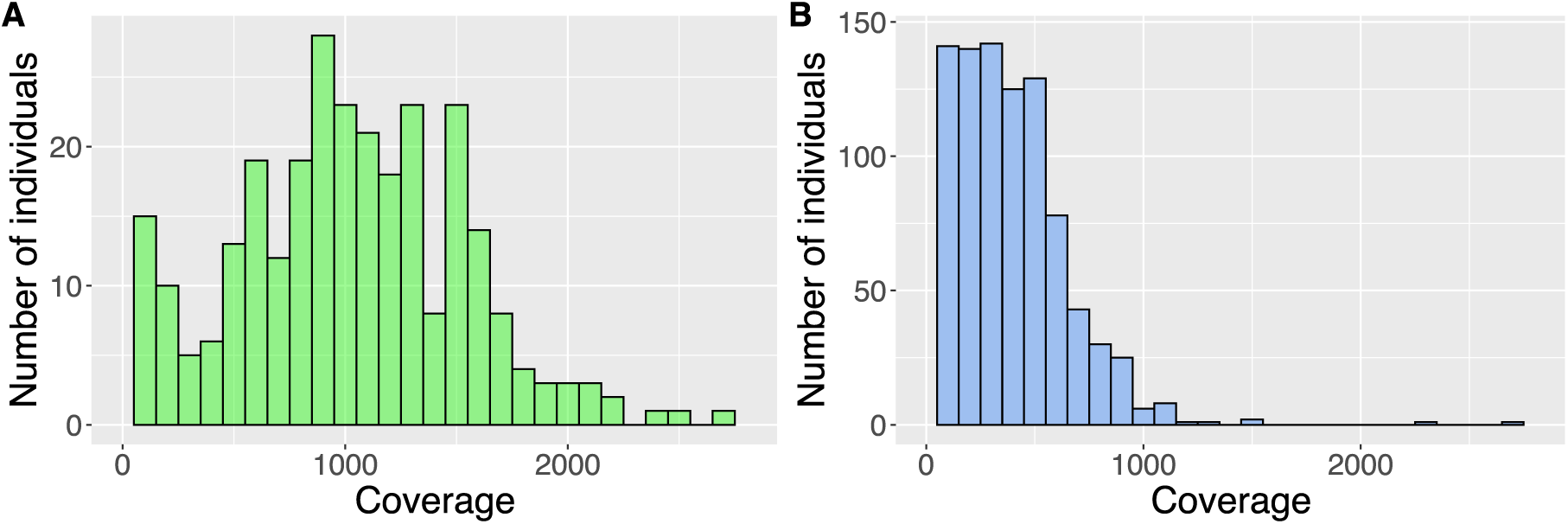
Histogram of mtDNA coverage for A) sequencing run 1, in which 292 samples were included. Average coverage is 994x, and B) for sequencing run 2, in which 1,024 samples were included. Average coverage is 340x.

**Supplementary Figure 4:**
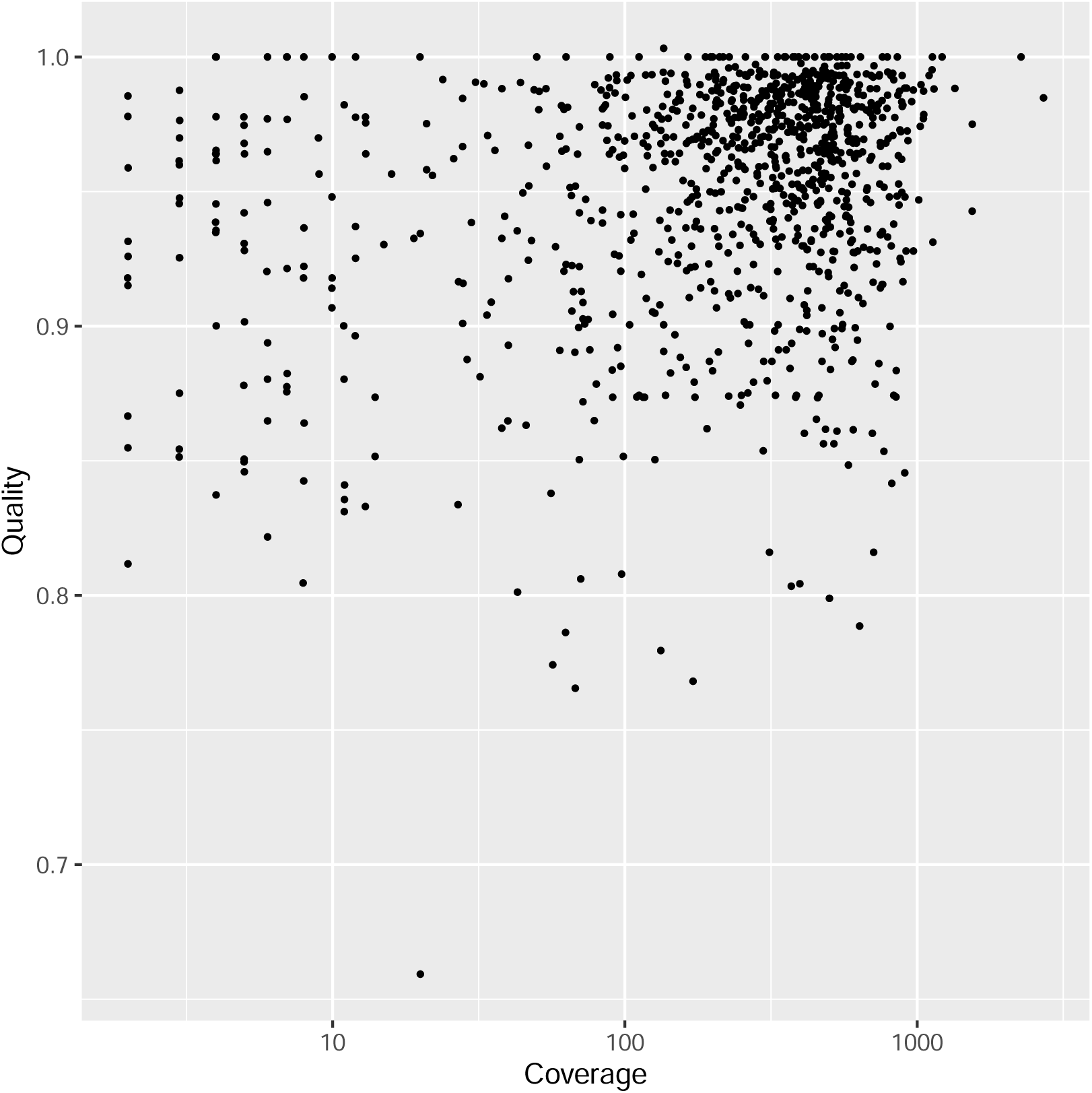
HaploGrep3 quality score of each sample plotted against its coverage of the mtDNA genome.

**Supplementary Figure 5:**
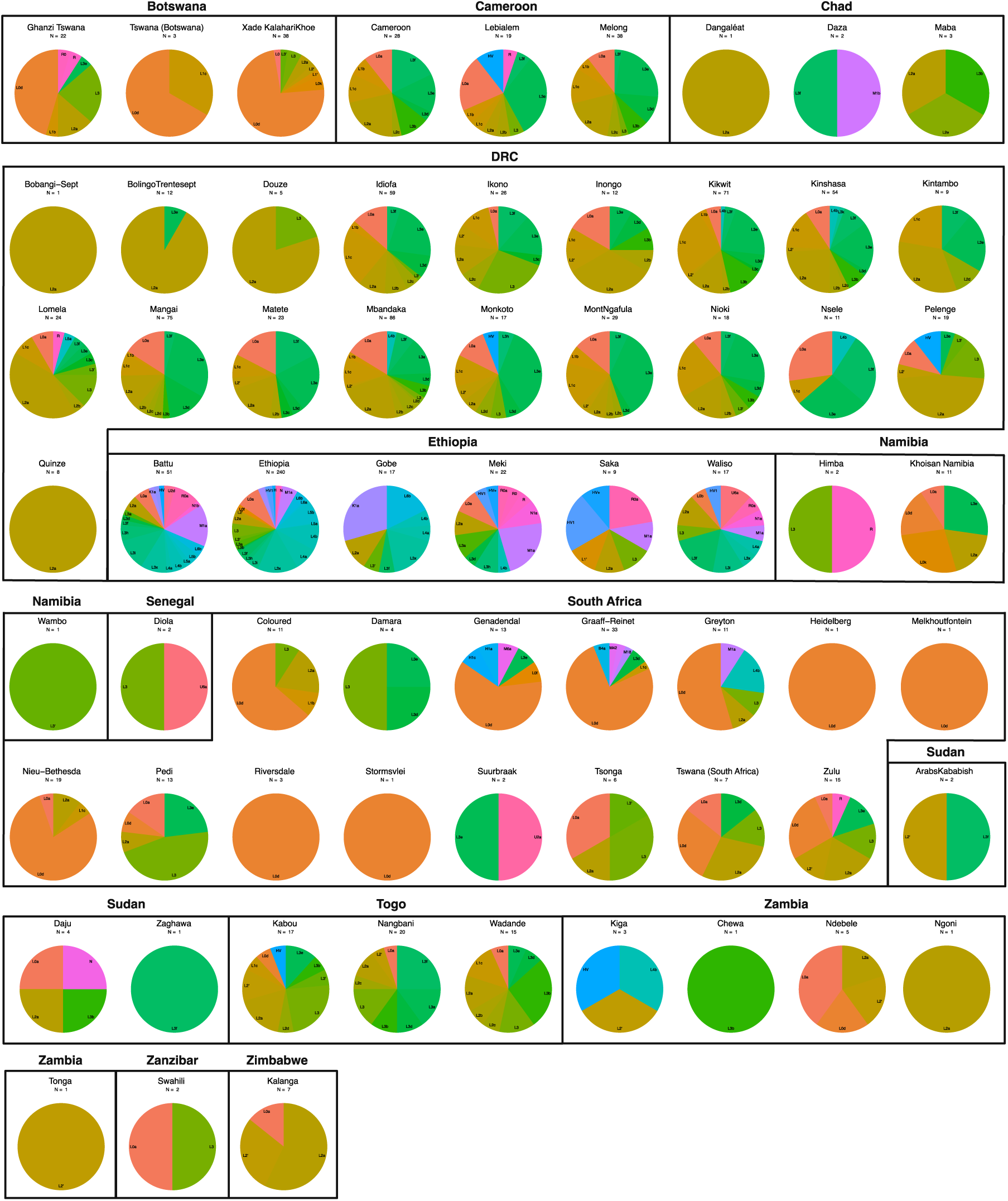
The mitochondrial haplogroups found among the studied individuals from the 66 sampling sites. Haplogroups were reduced to three-digit haplogroups. The number of individuals at each site is specified. Sites are grouped based on country. Colours do not correspond to ancestries reported in Supplementary Figure 6.

**Supplementary Figure 6:**
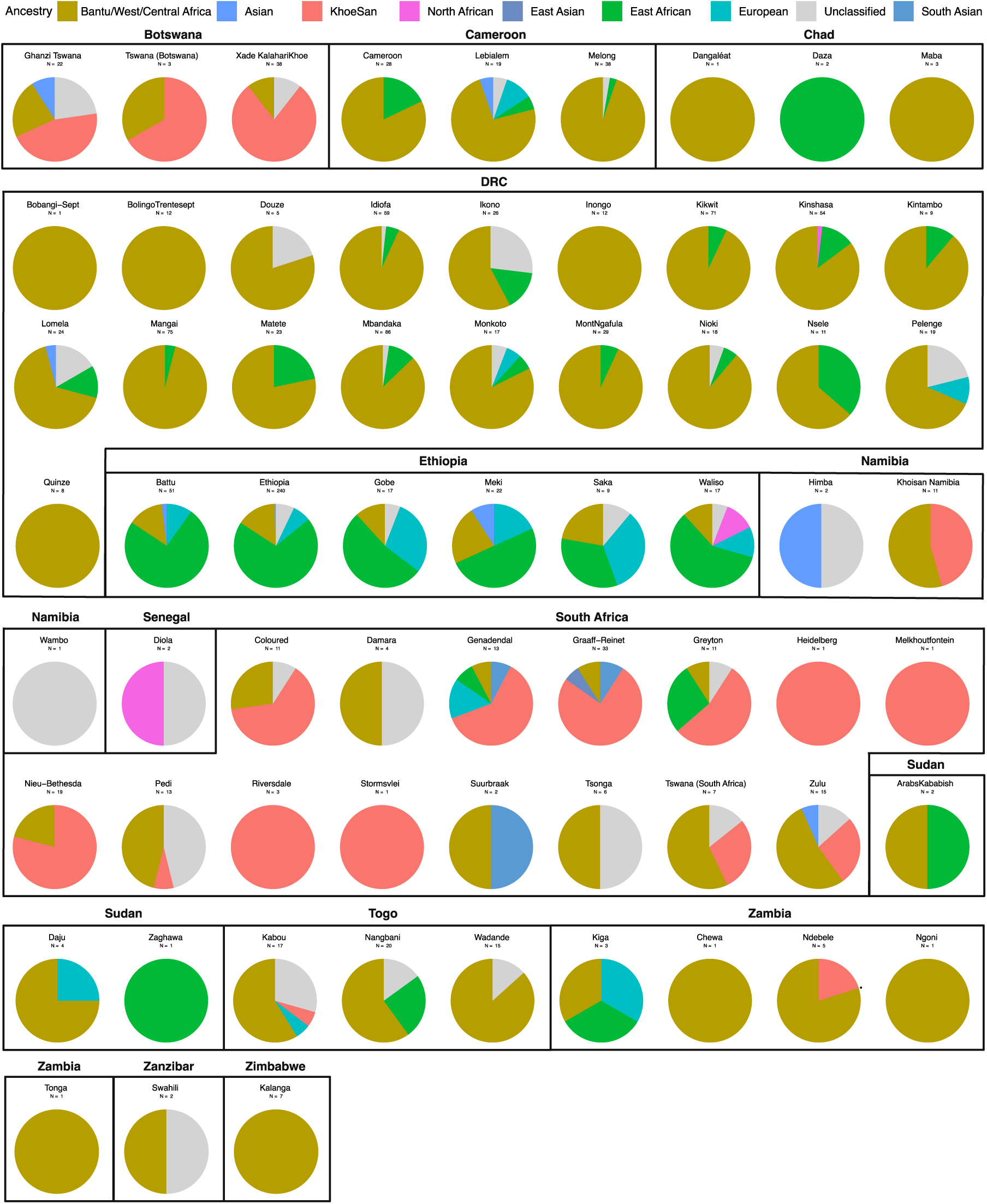
Maternal ancestries of individuals from 66 different sampling sites across 14 countries were determined by referencing maternal haplogroups in the literature (see Supplementary Table 4). The number of individuals at each site is specified. Sites are grouped based on country.

**Supplementary Figure 7:**
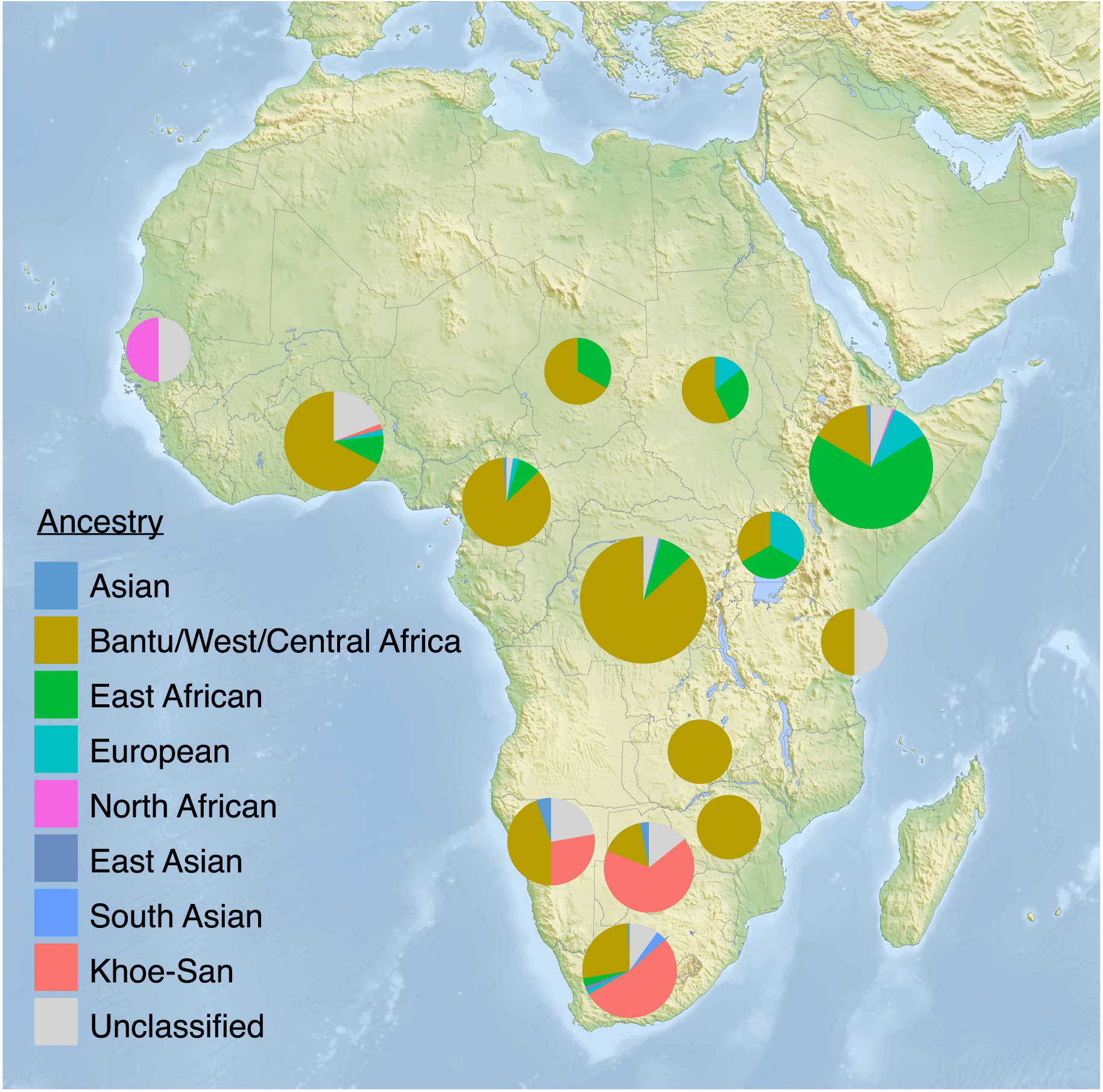
Maternal ancestries of individuals per country, visualized on a map. Ancestries were determined by referencing maternal haplogroups in the literature (see Supplementary Table 4).

**Supplementary Figure 8:**
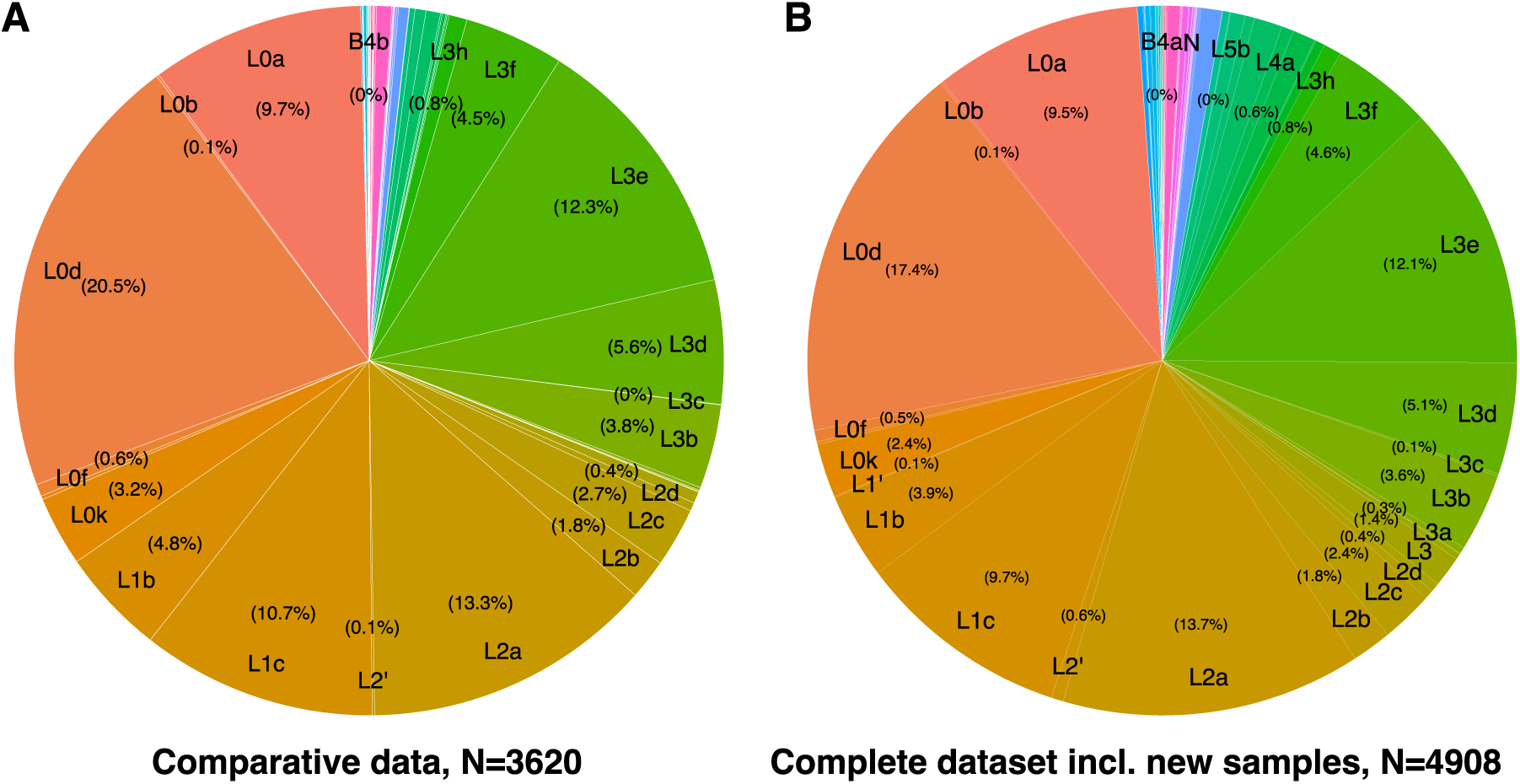
Distribution of the mitochondrial haplogroups of A) the 3,620 individuals in the comparative dataset, and B) the 4,908 individuals in the complete dataset. This includes the newly sequenced 1,288 individuals. The distribution of mitochondrial haplogroups shown here is heavily influenced by sampling bias (a focus on Khoe-San groups in published literature can probably explain why L0d is the most frequent mitochondrial haplogroup in our dataset). This is therefore not necessarily a reflection of the “real” frequency of mitochondrial haplogroups across the African continent.

**Supplementary Figure 9:**
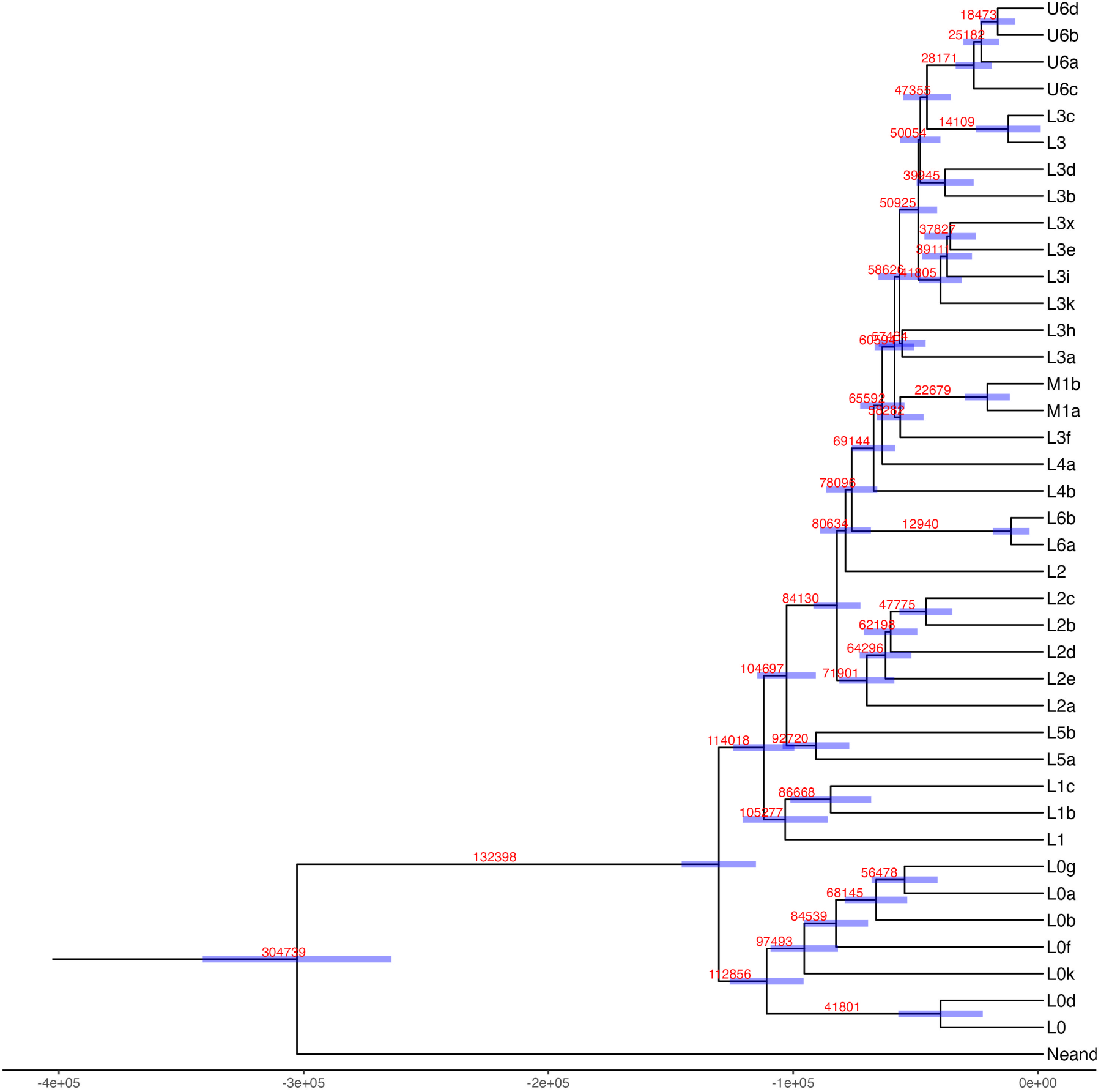
The Bayesian tree topology of African mitochondrial haplogroups with Time to Most Recent Common Ancestor (TMRCA) information. Blue bars indicate the 95% confidence intervals of common ancestor heights; in red the mean common ancestor heights are shown.

**Supplementary Figure 10:**
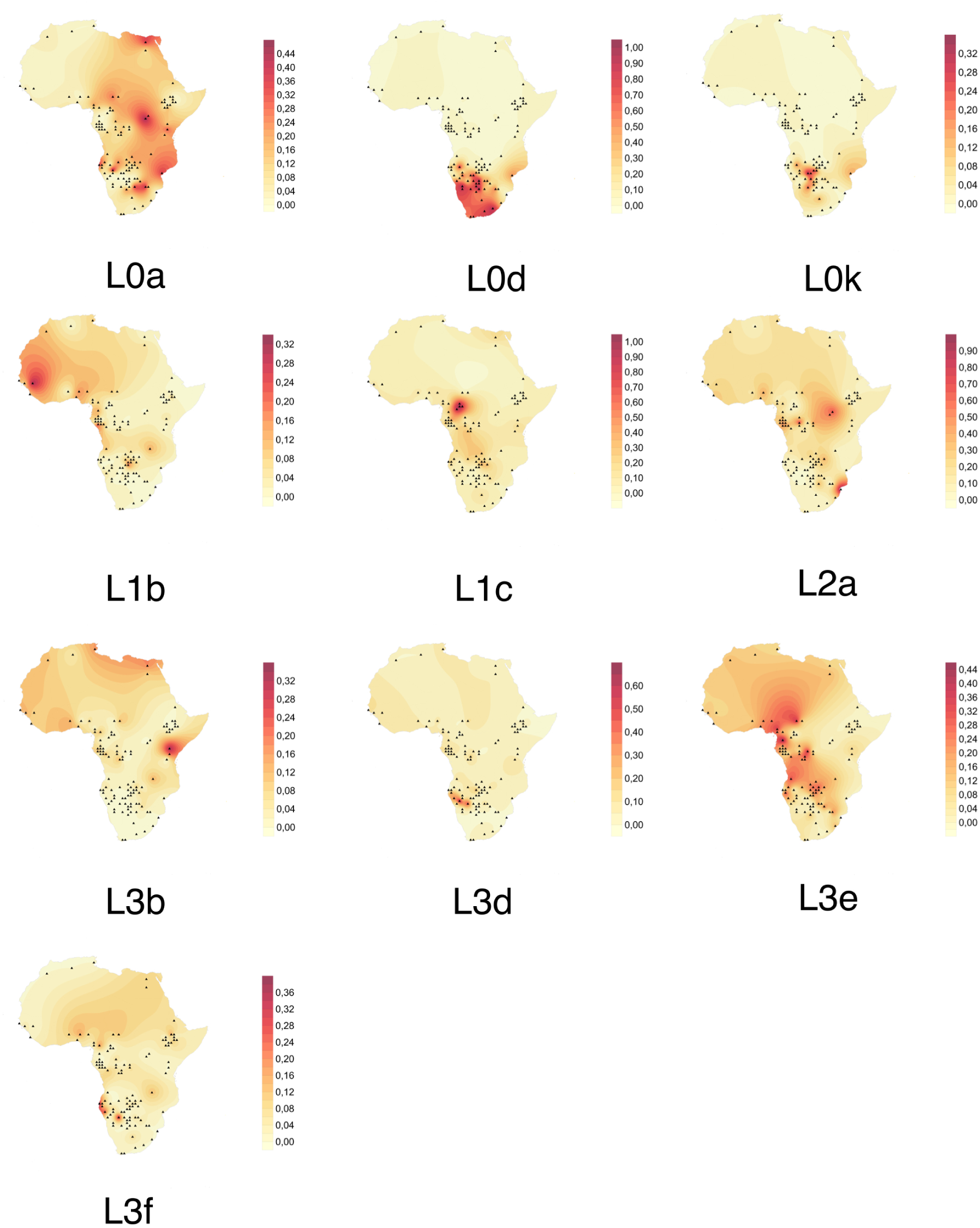
Surfer maps of spatial distribution of the haplogroup frequencies for the 10 haplogroups most commonly found in our dataset. The Kriging method was applied. Note that the intensities of the colours are not comparable between the different subfigures.

**Supplementary Figure 11:**
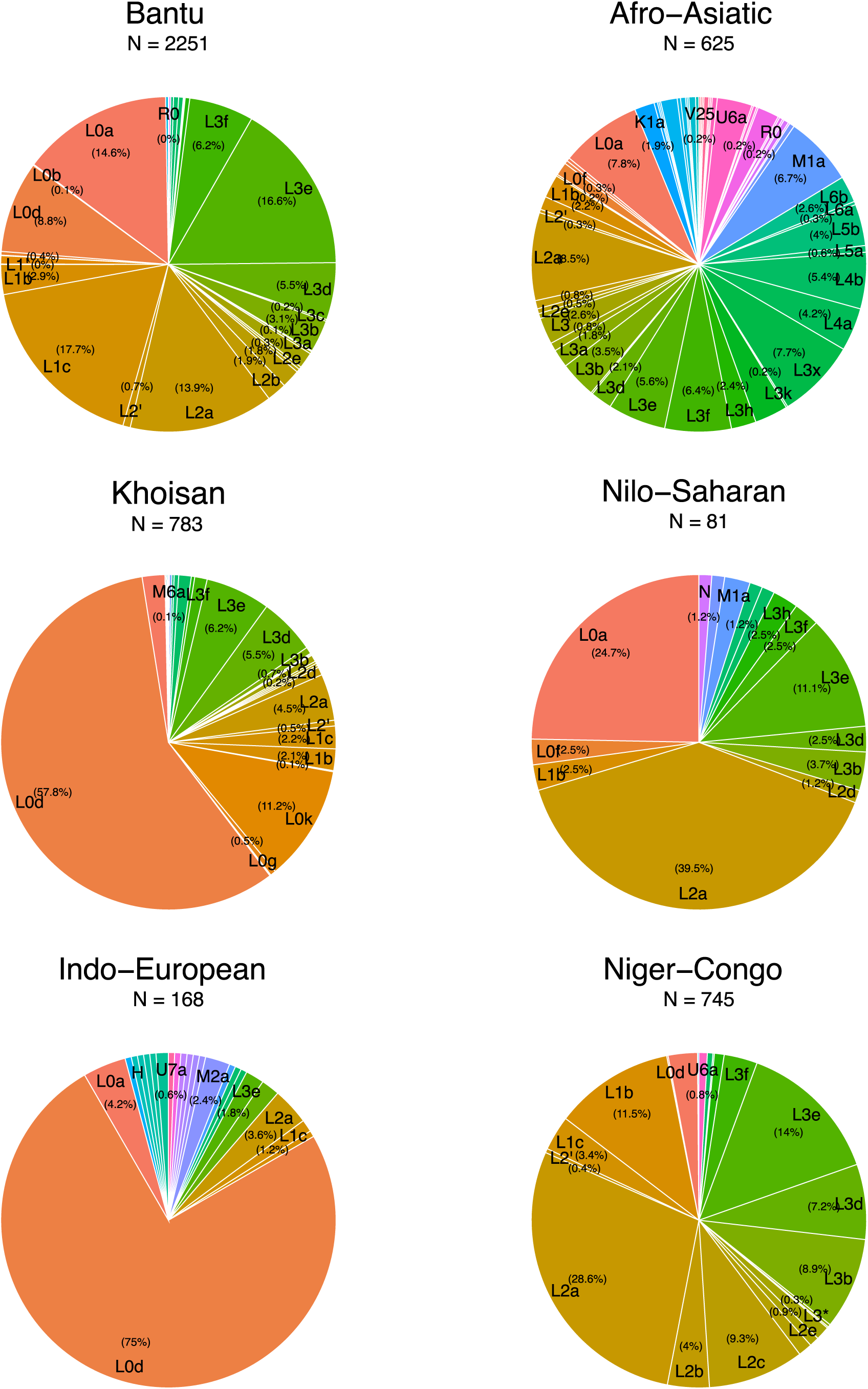
Distribution of the mitochondrial haplogroups among the different language groups.

**Supplementary Figure 12:**
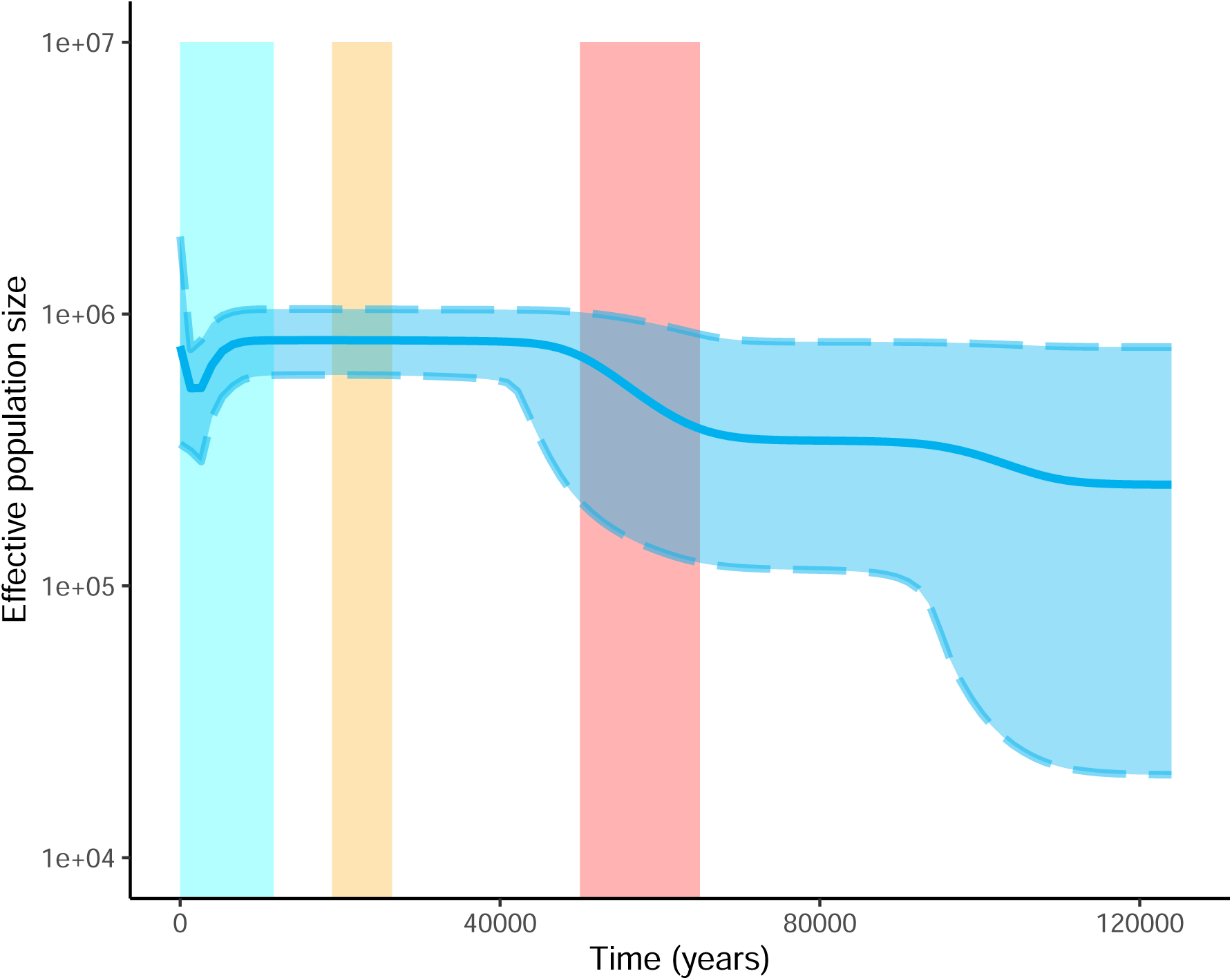
Bayesian Skyline Plots showing the variation of female *N*_e_ through time for all Khoisan speakers. The bold middle line represents the mean estimates and the two thinner lines surrounding the bold line represent the 95% highest posterior density (HPD) intervals. The red area denotes the time of the out-of-Africa migration (65–50 kya), the yellow area the Last Glacial Maximum (LGM)(26.5–19 kya), and the blue area the Holocene (last 11.7 ky). For the generation of this plot, 783 individuals were included.

**Supplementary Figure 13:**
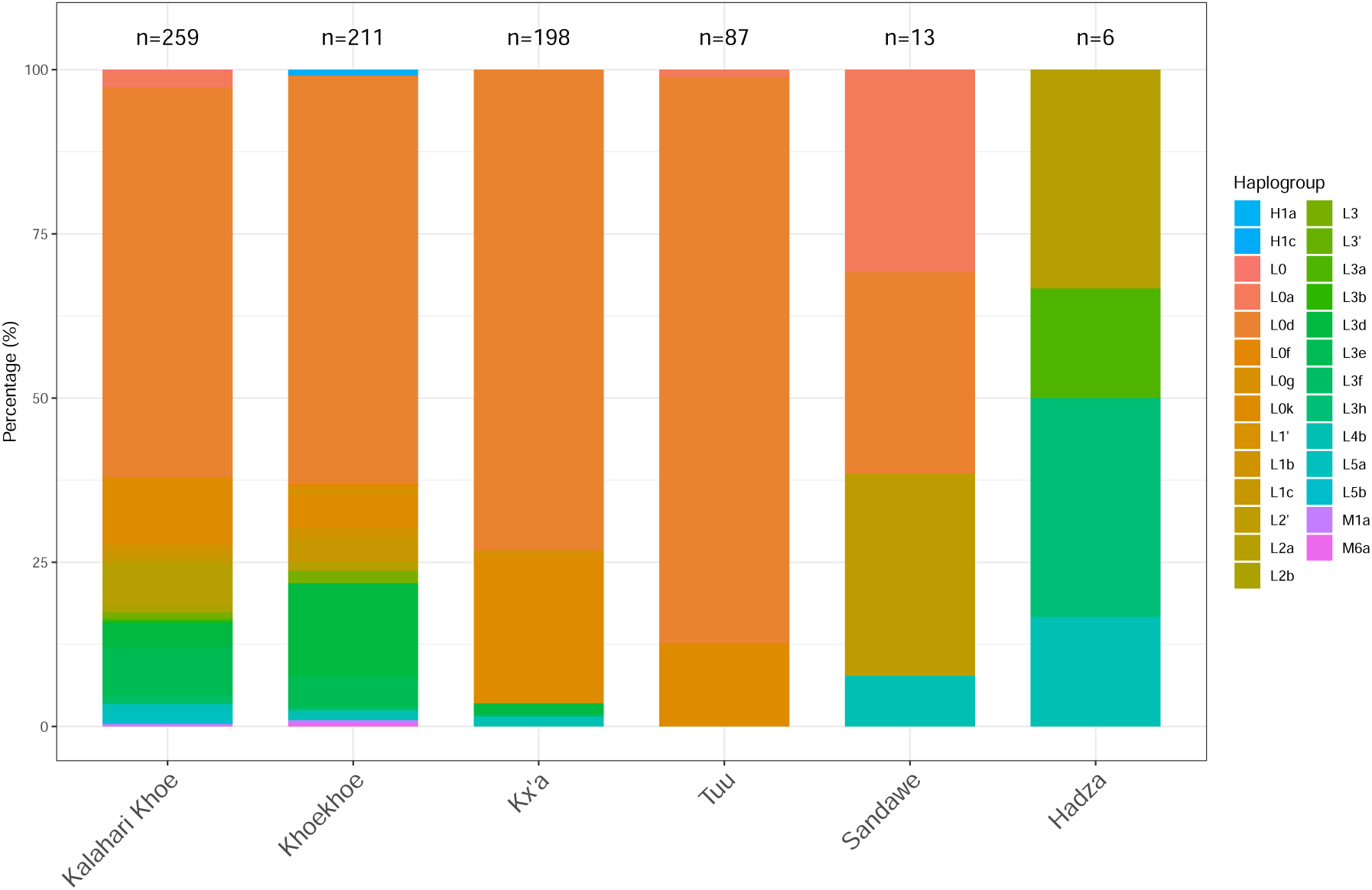
Haplogroup distribution among the six Khoisan languages. Number of individuals in each group is denoted at the top of each bar.

**Supplementary Figure 14:**
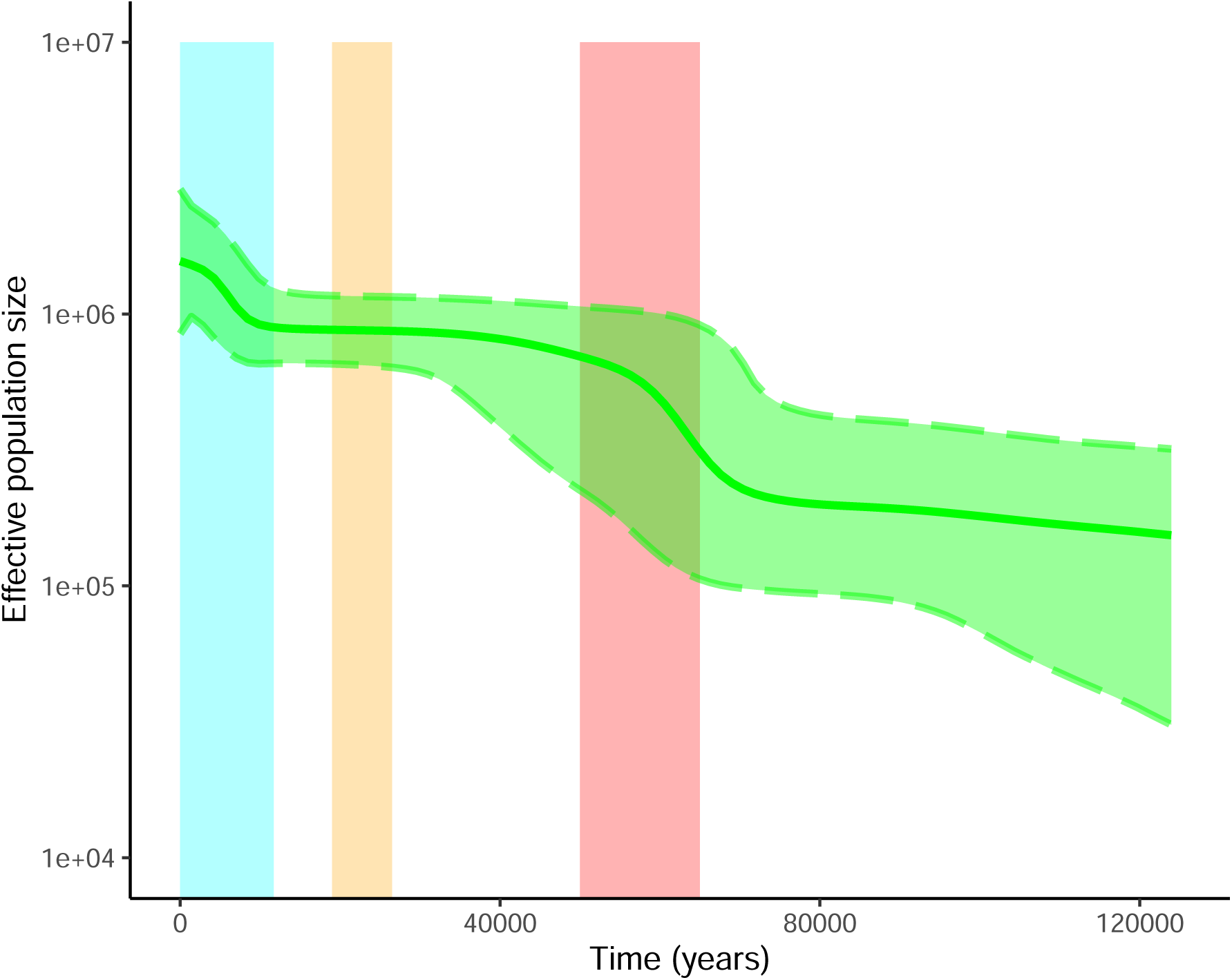
Bayesian Skyline Plots showing the variation of *N*_e_ through time for all Afrikaans speakers in the dataset. The bold middle line represents the mean estimates and the two thinner lines surrounding the bold line represent the 95% highest posterior density (HPD) intervals. The red area denotes the time of the out-of-Africa migration (65–50 kya), the yellow area the Last Glacial Maximum (LGM)(26.5–19 kya), and the blue the Holocene (last 11.7 ky). For the generation of this plot, 211 individuals were included.

**Supplementary Figure 15:**
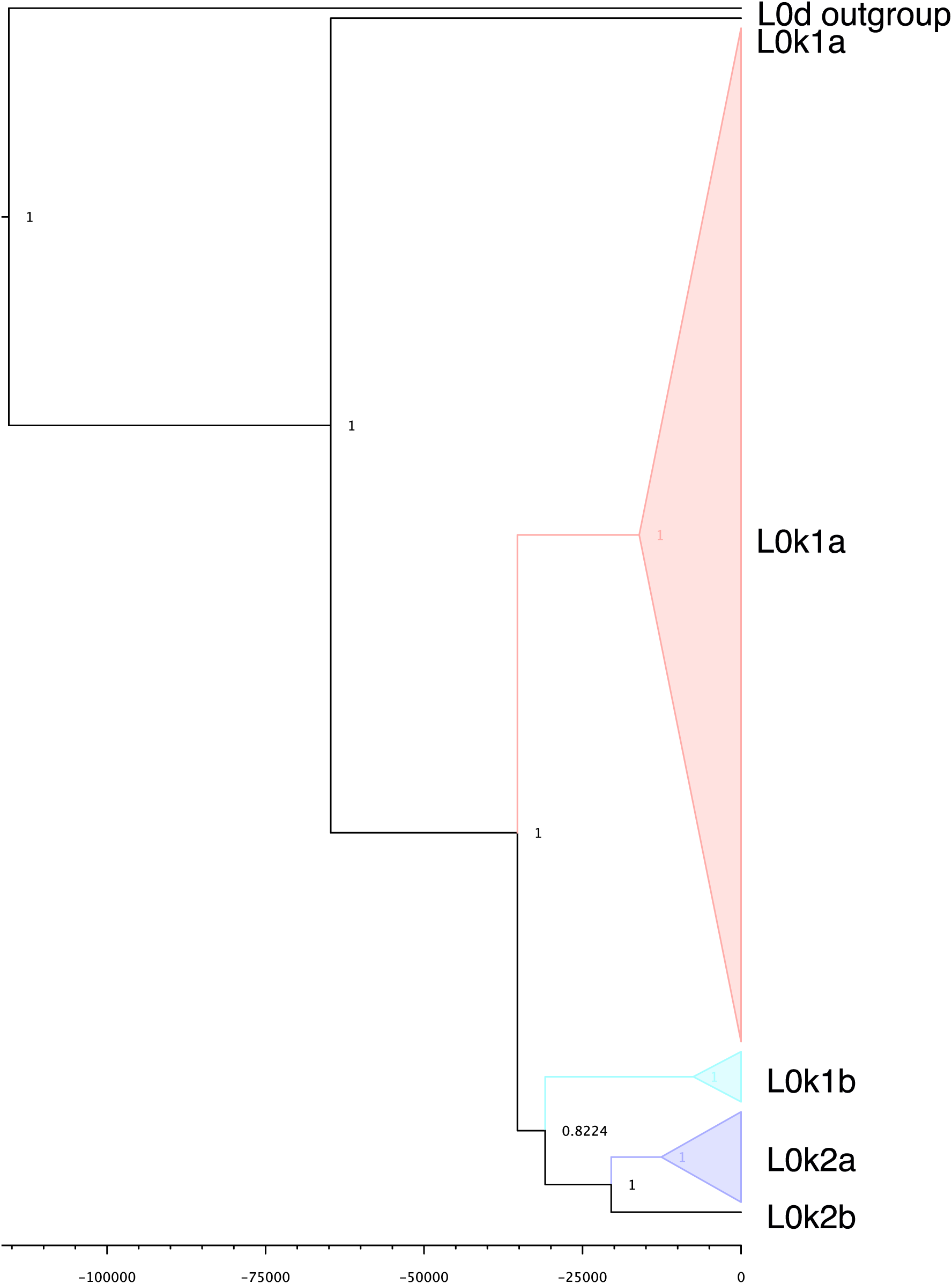
A maximum clade credibility tree for all the sequences in the dataset belonging to mitochondrial haplogroup L0k. Node heights represent common ancestor heights. Posterior probability is given at the nodes. On the X-axis, the years in the past are denoted.

**Supplementary Figure 16:**
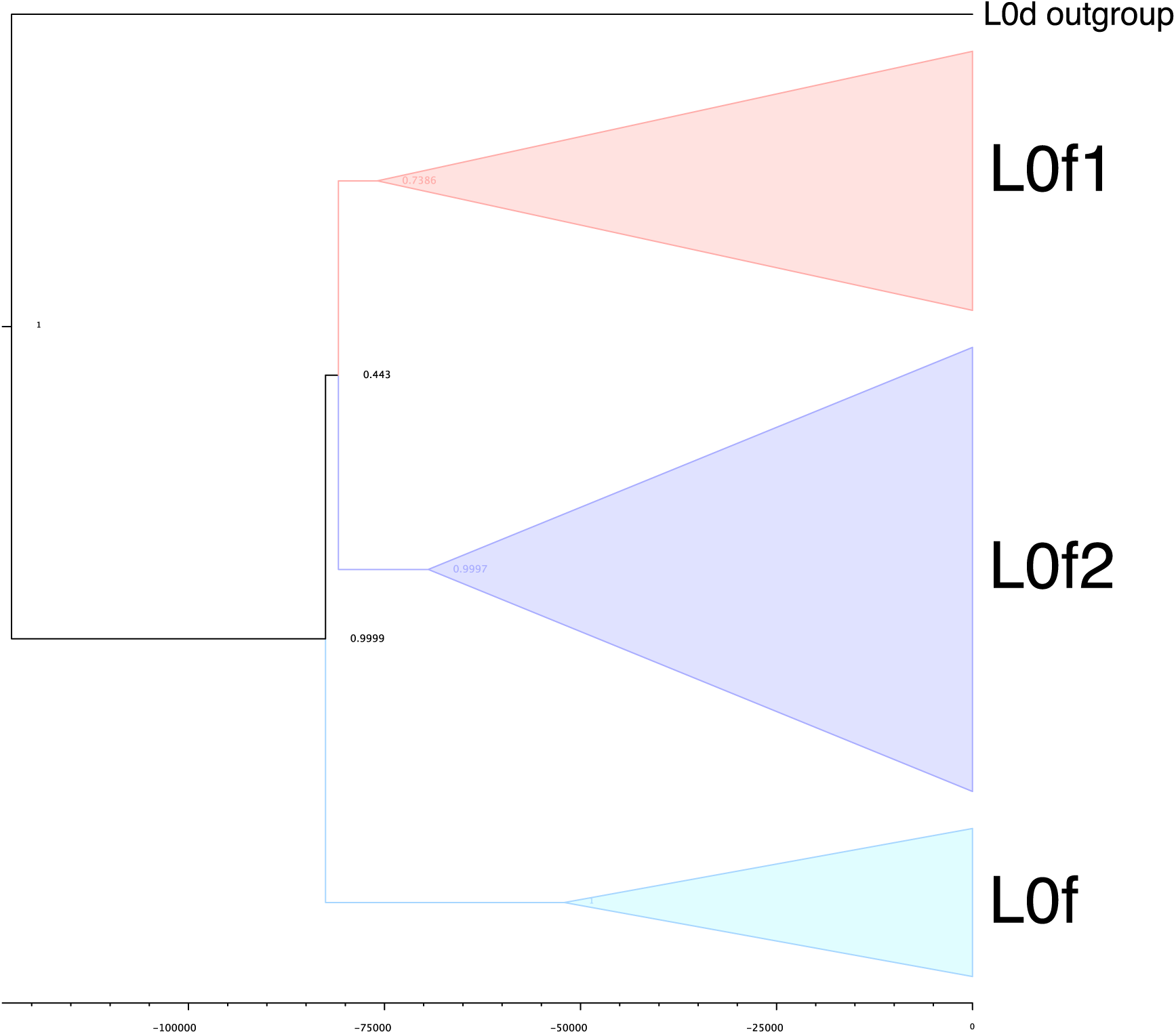
A maximum clade credibility tree for all the sequences in the dataset belonging to mitochondrial haplogroup L0f. Node heights represent common ancestor heights. Posterior probability is given at the nodes. On the X-axis, the years in the past are denoted.

**Supplementary Figure 17:**
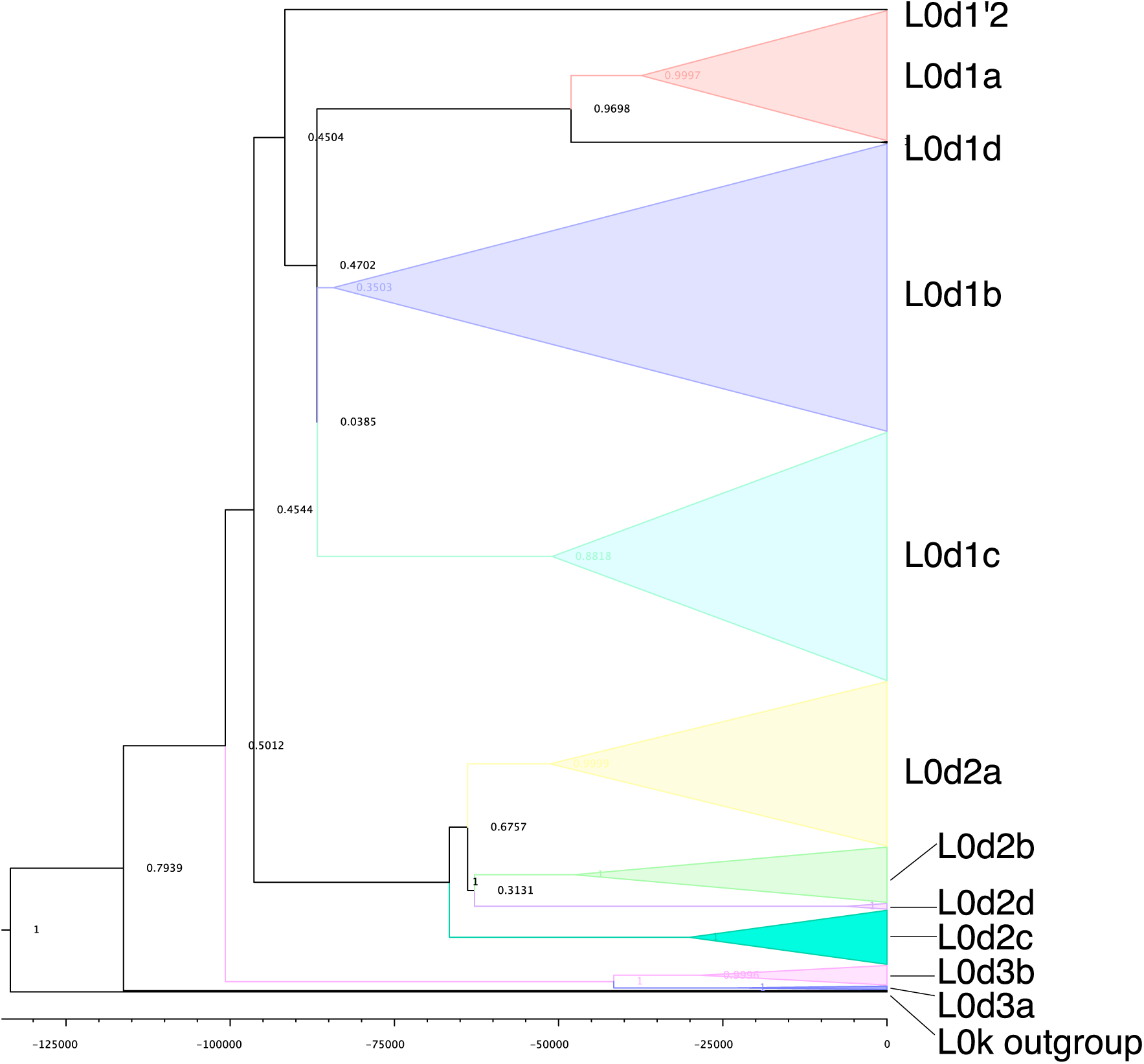
A maximum clade credibility tree for all the sequences in the dataset belonging to mitochondrial haplogroup L0d. Node heights represent common ancestor heights. Posterior probability is given at the nodes. On the X-axis, the years in the past are denoted.

**Supplementary Figure 18:**
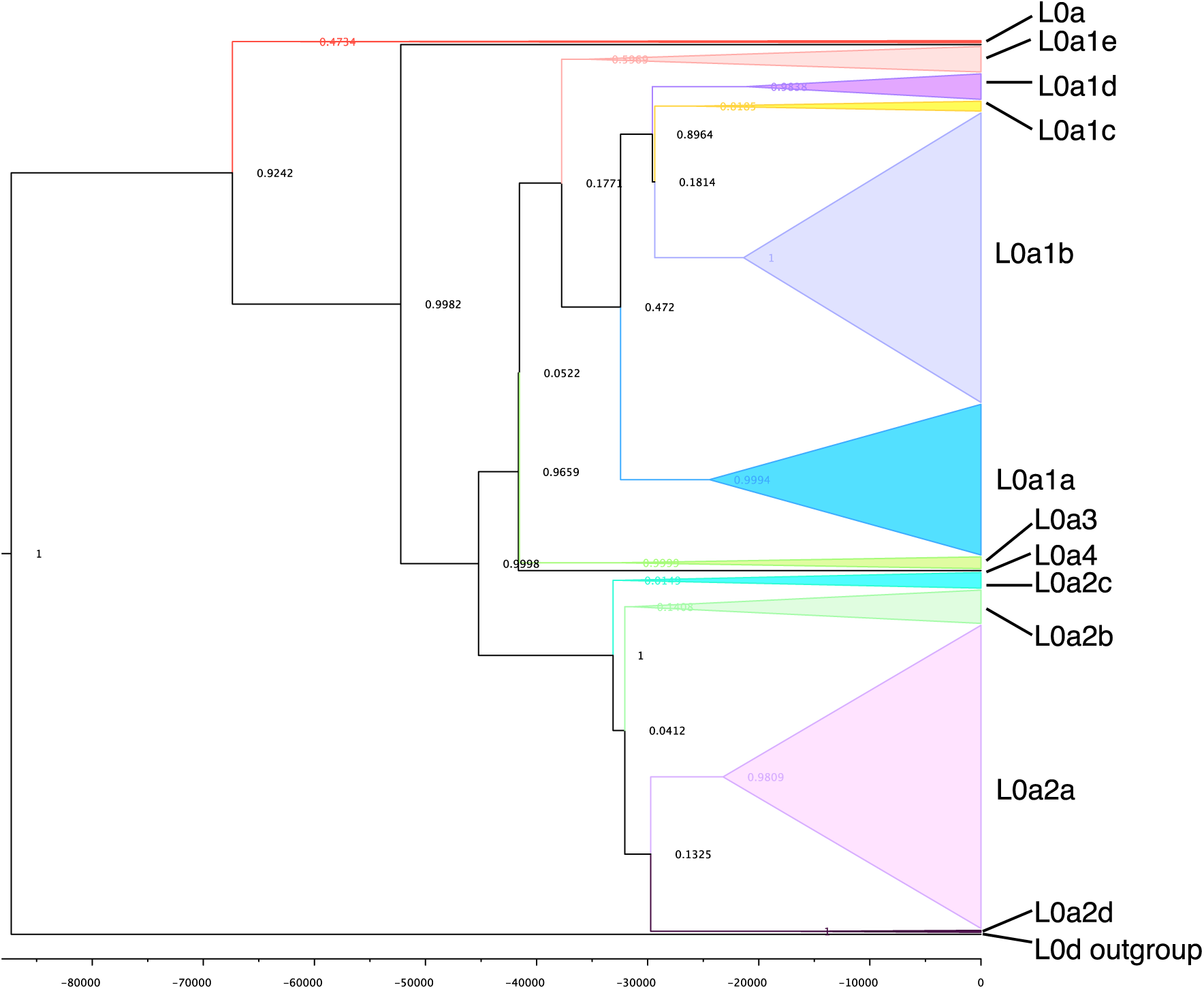
A maximum clade credibility tree for all the sequences in the dataset belonging to mitochondrial haplogroup L0a. Node heights represent common ancestor heights. Posterior probability is given at the nodes. On the X-axis, the years in the past are denoted.

**Supplementary Figure 19:**
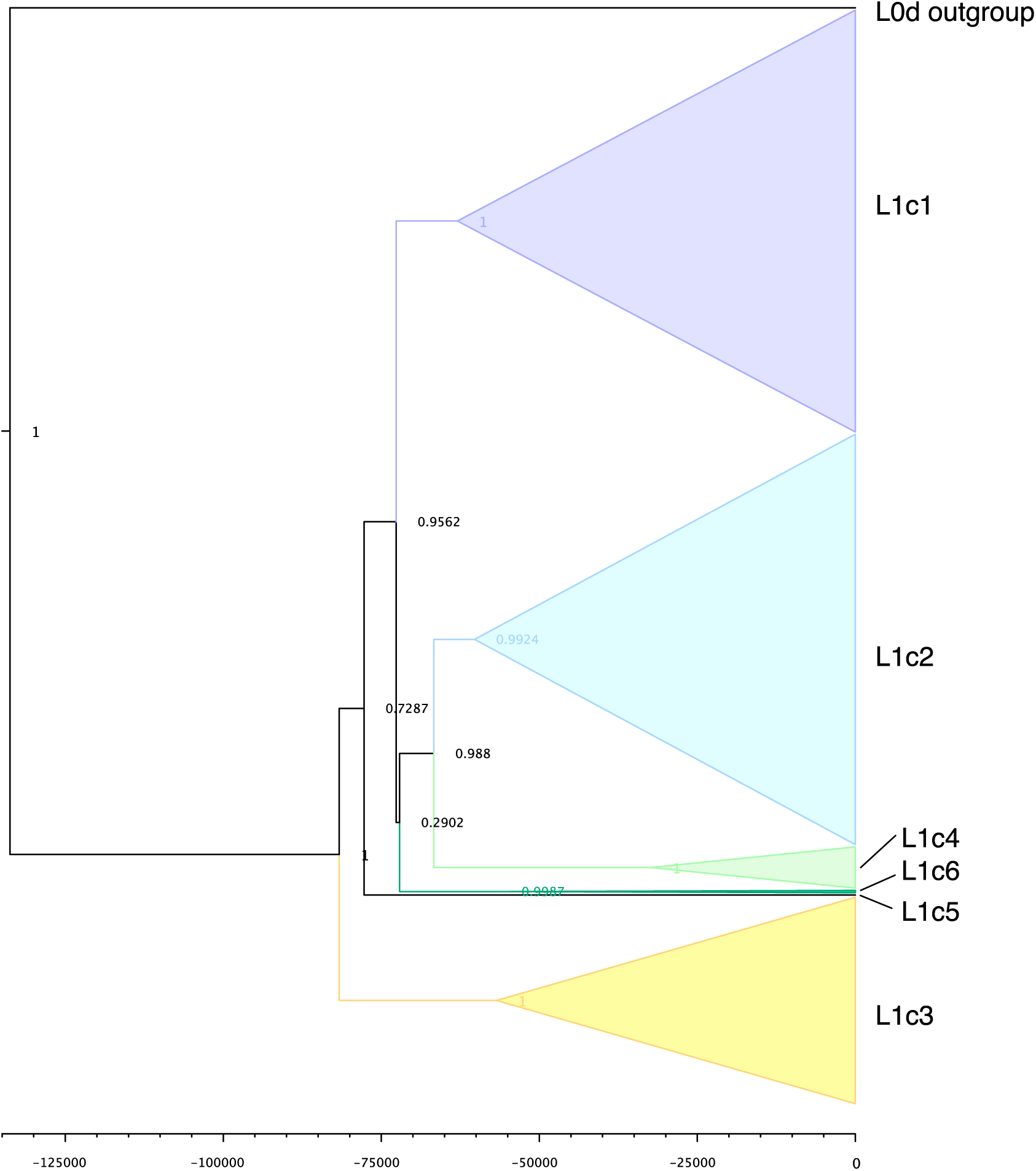
A maximum clade credibility tree for all the sequences in the dataset belonging to mitochondrial haplogroup L1c. Node heights represent common ancestor heights. Posterior probability is given at the nodes. On the X-axis, the years in the past are denoted.

**Supplementary Figure 20:**
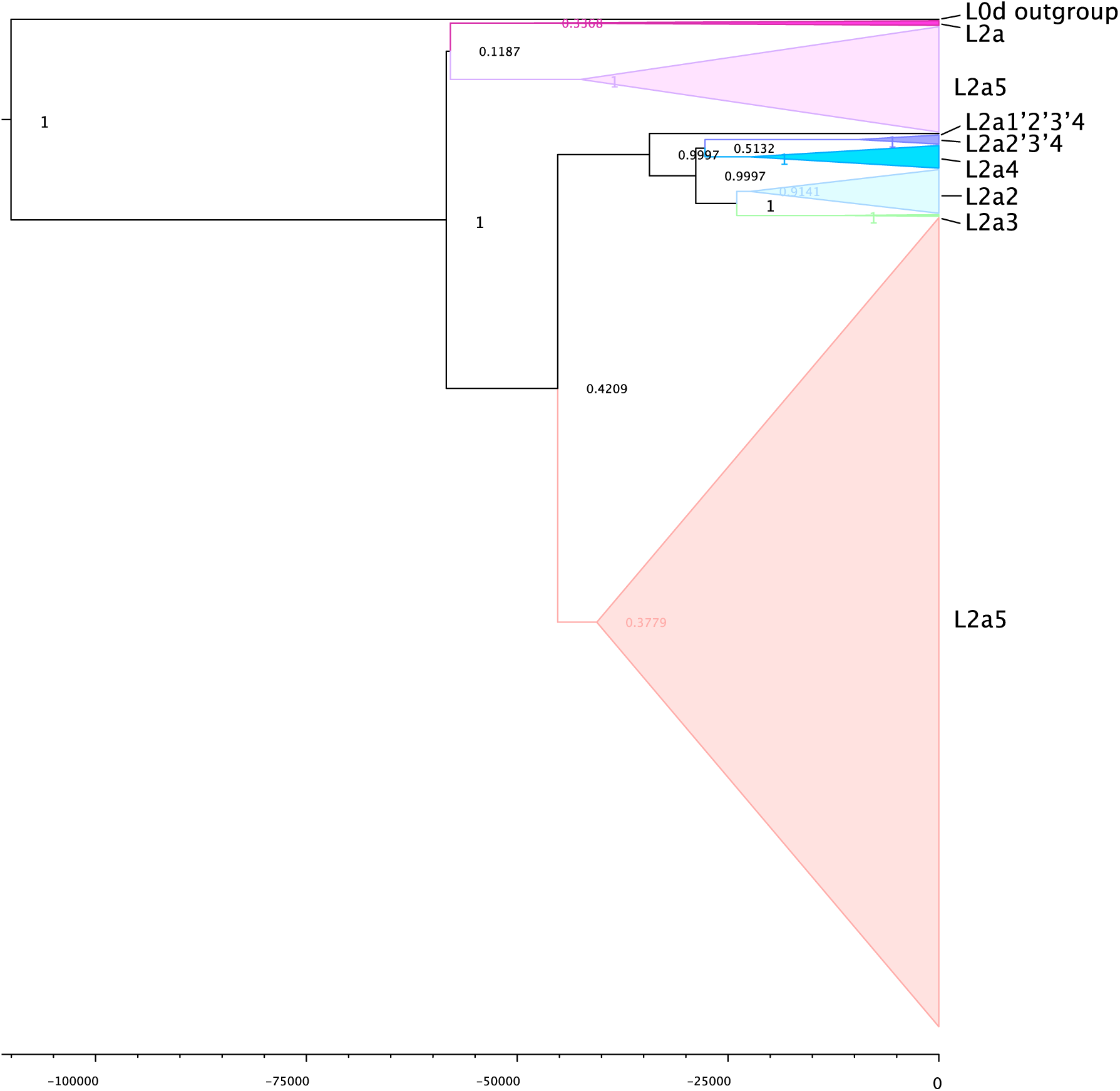
A maximum clade credibility tree for all the sequences in the dataset belonging to mitochondrial haplogroup L2a. Node heights represent common ancestor heights. Posterior probability is given at the nodes. On the X-axis, the years in the past are denoted.

**Supplementary Figure 21:**
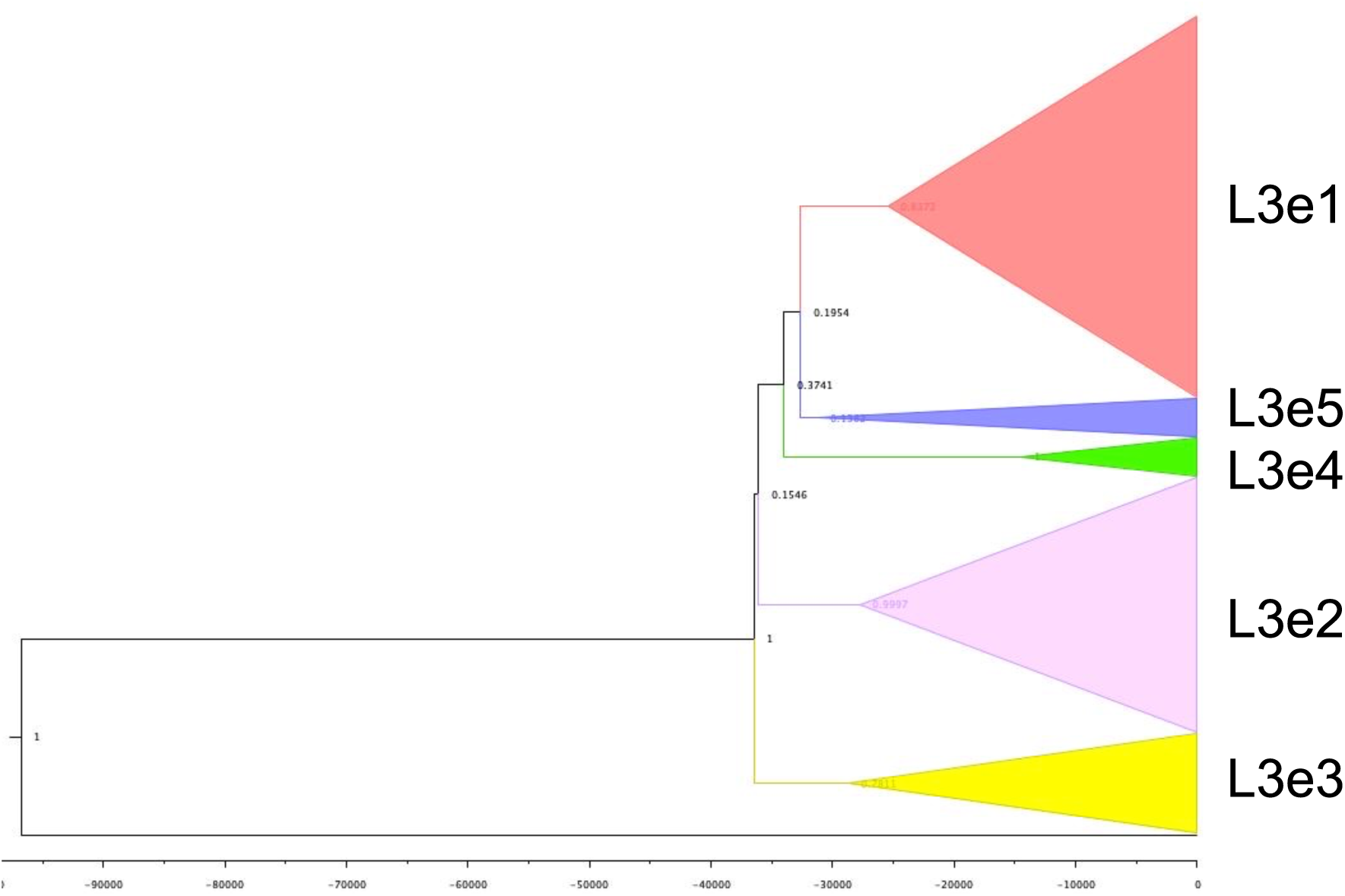
A maximum clade credibility tree for all the sequences in the dataset belonging to mitochondrial haplogroup L3e. Node heights represent common ancestor heights. Posterior probability is given at the nodes. On the X-axis, the years in the past are denoted.

**Supplementary Figure 22:**
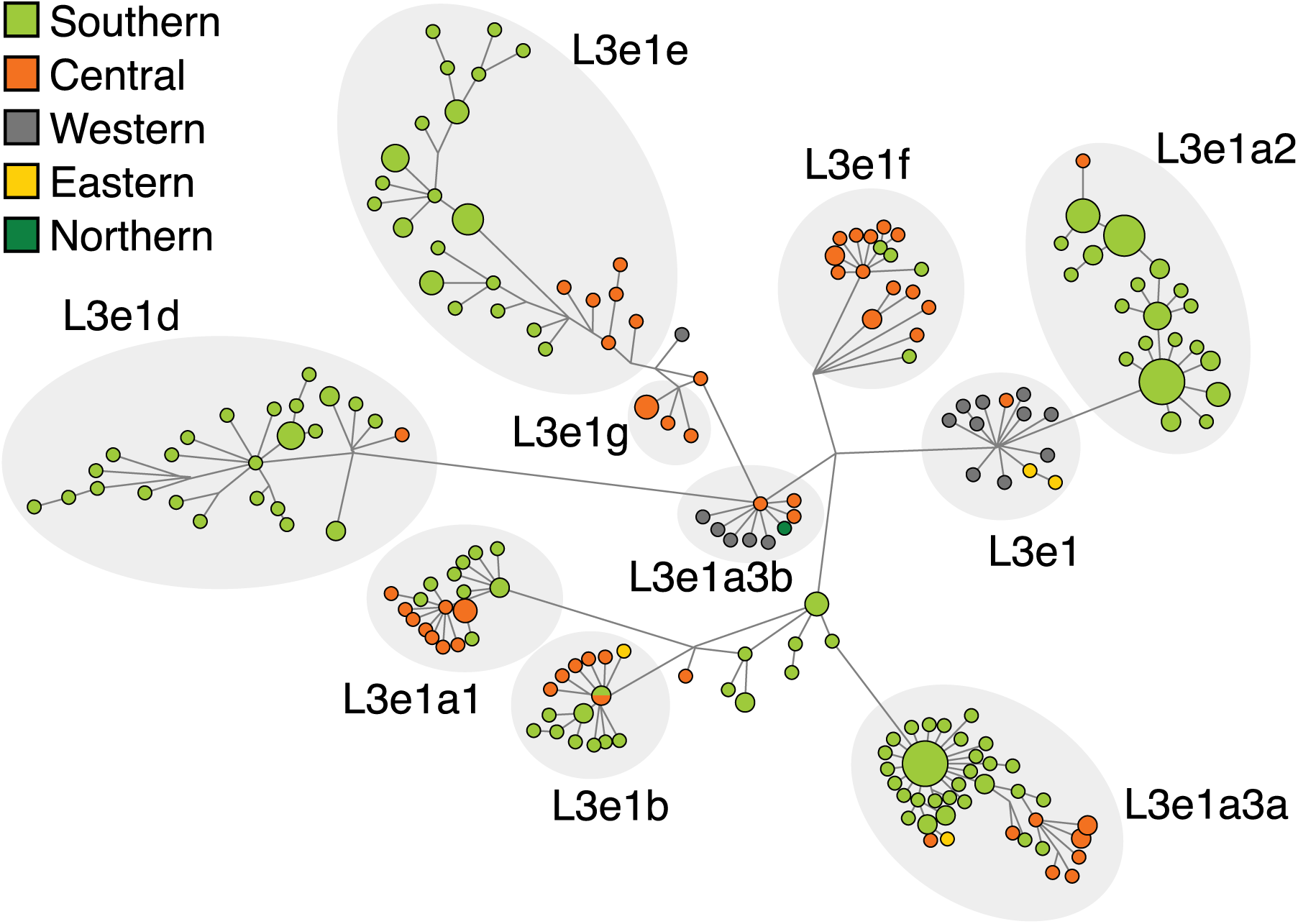
Median joining network of all sequences belonging to mitochondrial haplogroup L3e1.

**Supplementary Figure 23:**
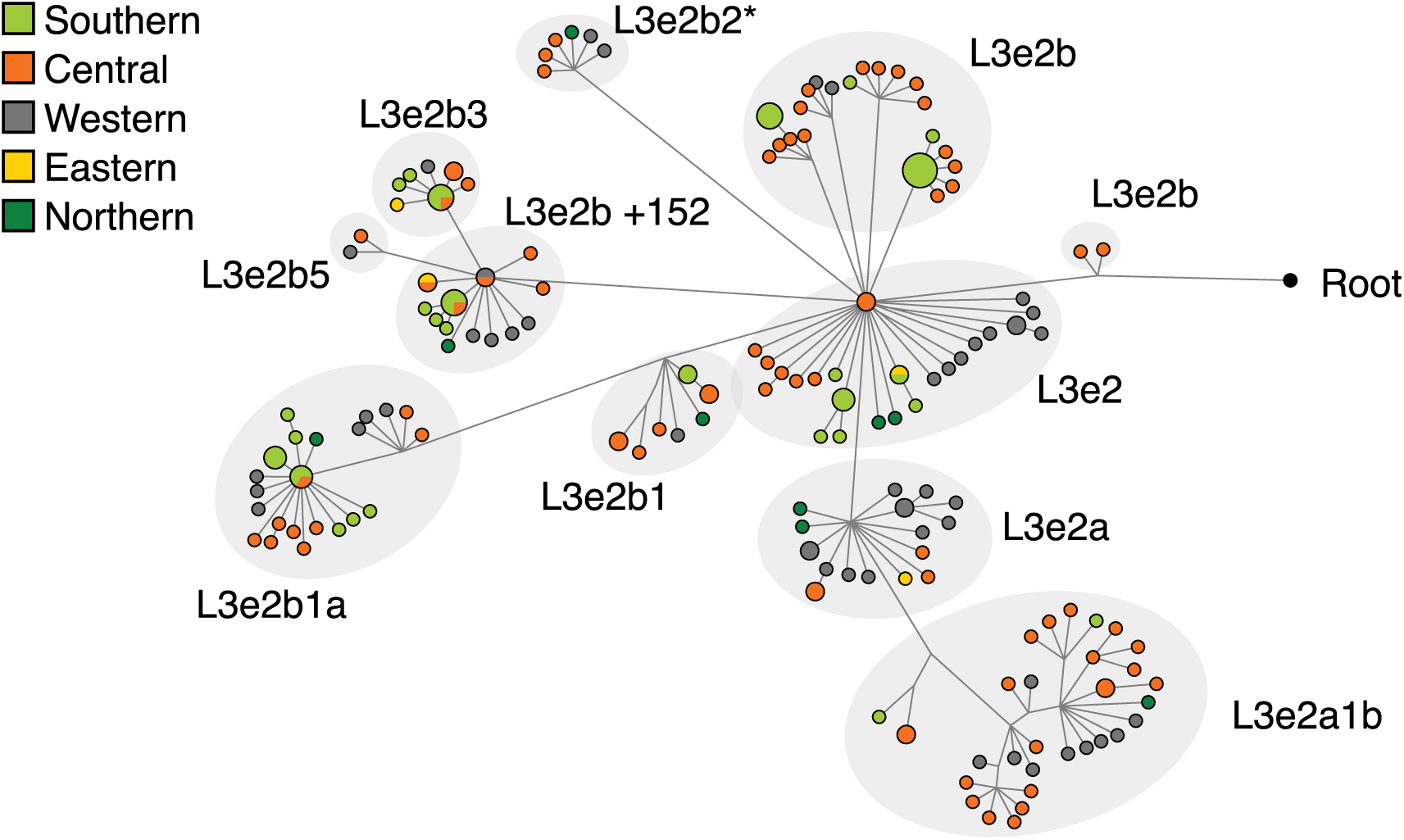
Median joining network of all sequences belonging to mitochondrial haplogroup L3e2.

**Supplementary Figure 24:**
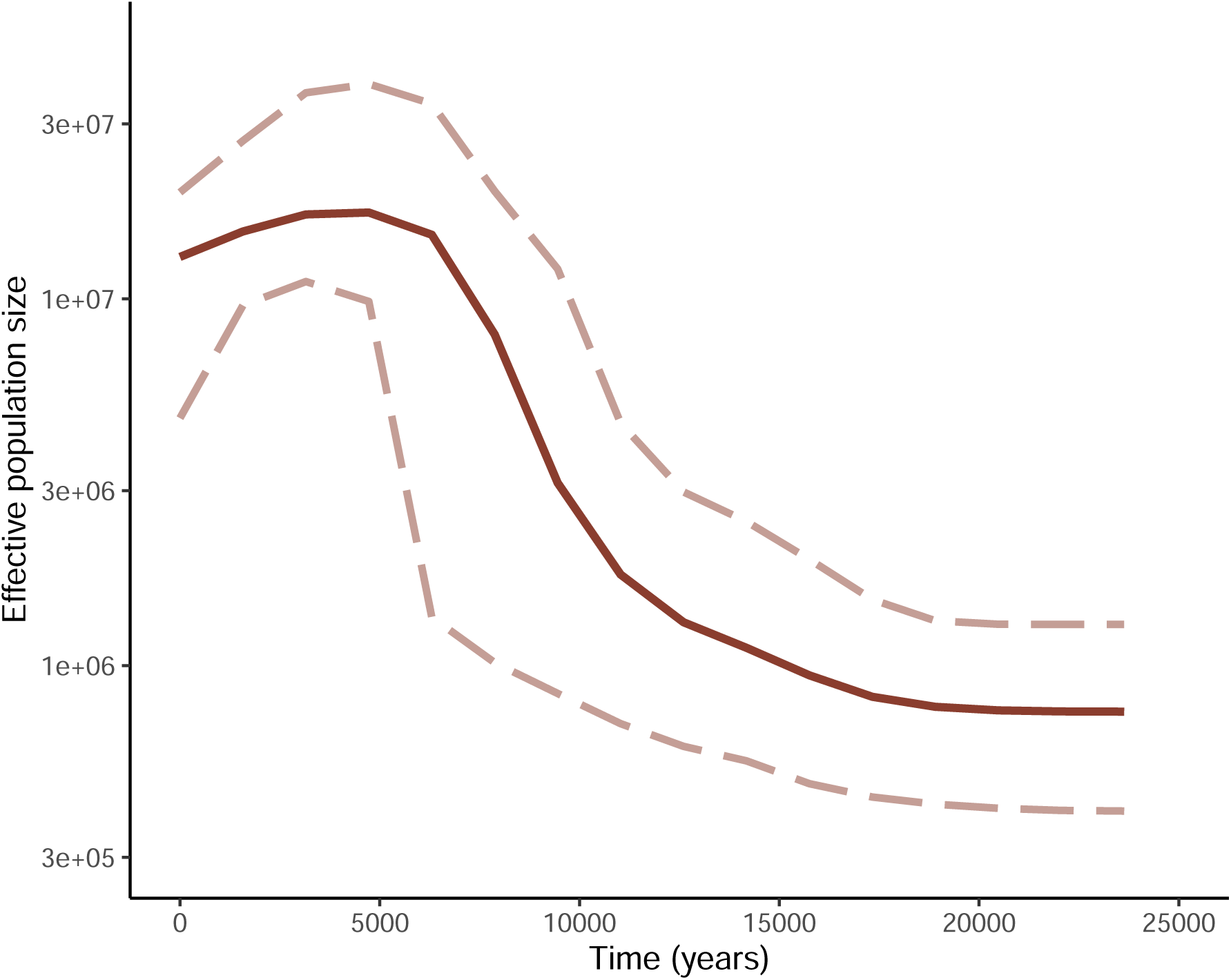
Bayesian Skyline Plots showing the variation of *N*_e_ through time for Niger-Congo and Mande speakers, excluding Bantu speakers.

**Supplementary Figure 25:**
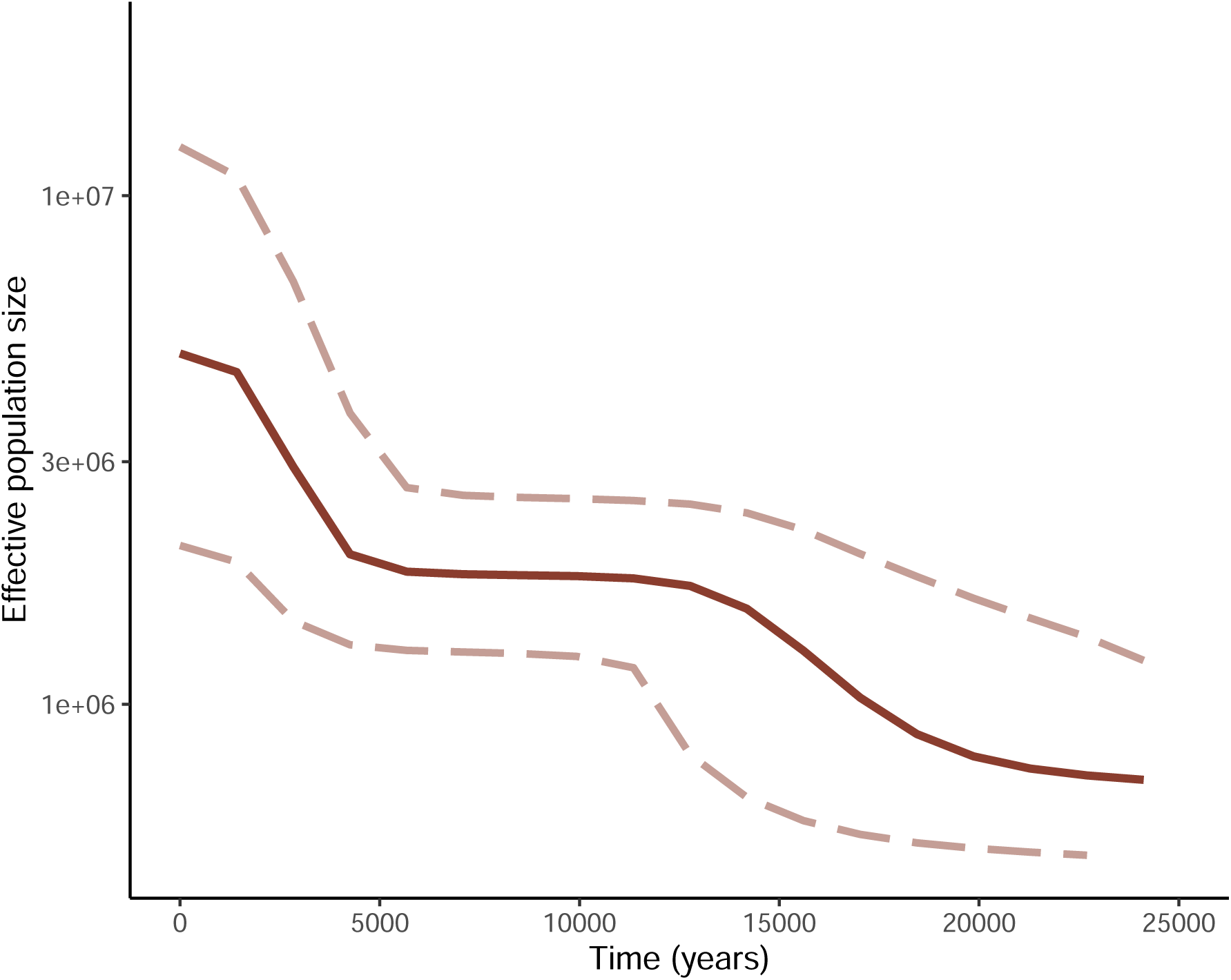
Bayesian Skyline Plots showing the variation of *N*_e_ through time for Niger-Congo and Mande speakers, including Bantu speakers.

**Supplementary Figure 26:**
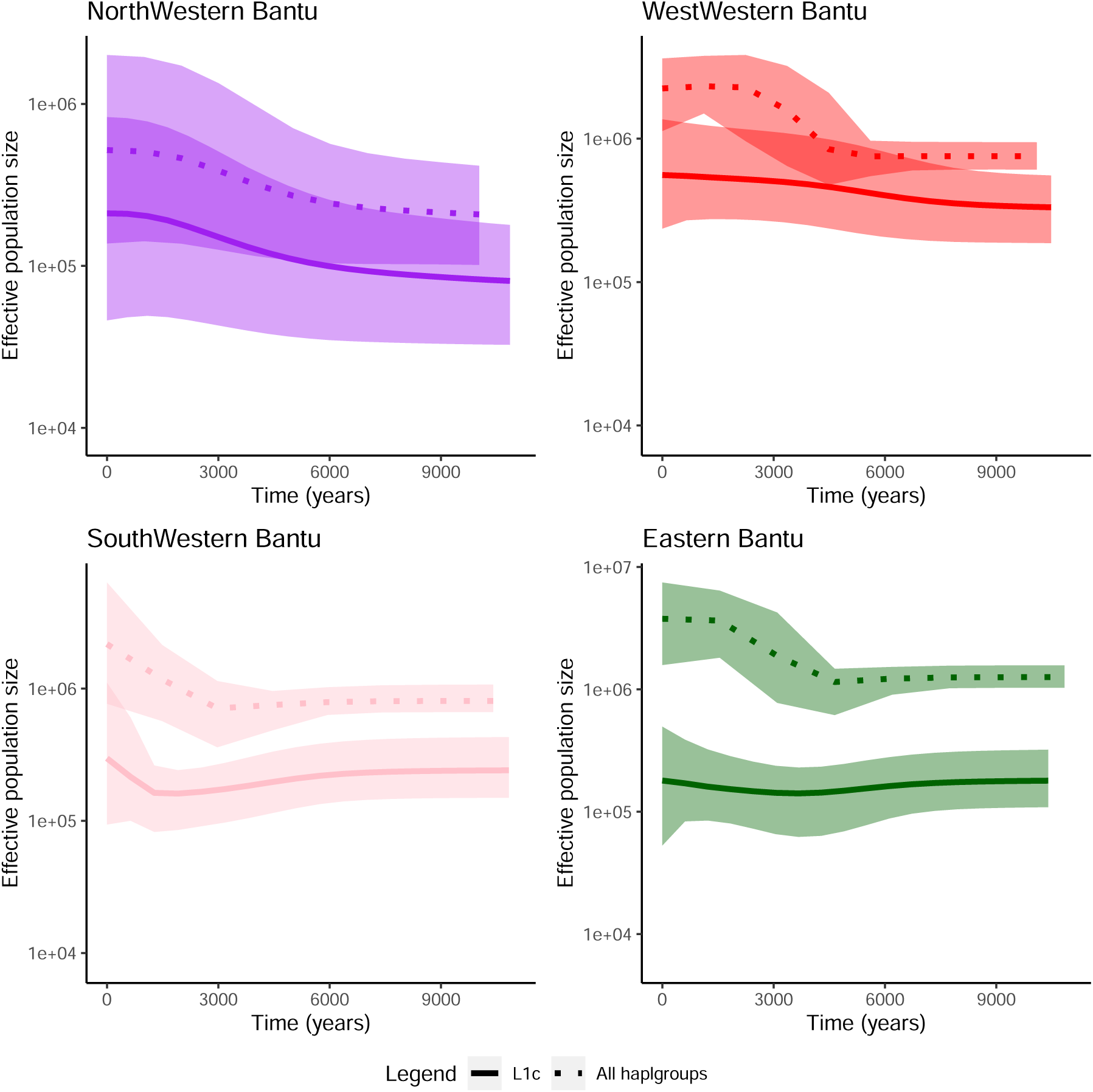
Bayesian Skyline Plots showing the variation of *N*_e_ through time estimated from L1c carriers of separate Bantu speaker groups.

**Supplementary Figure 27:**
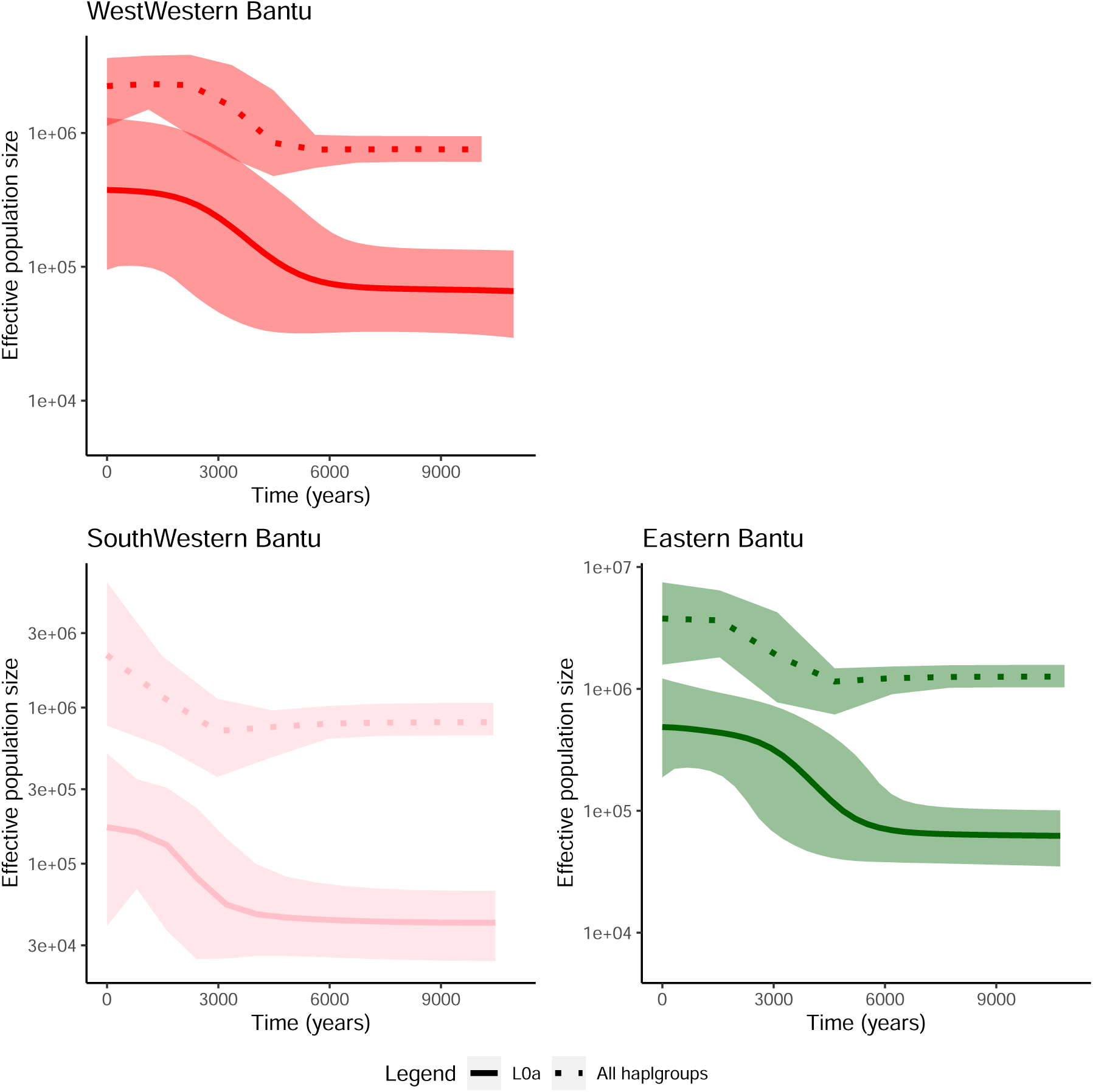
Bayesian Skyline Plots showing the variation of *N*_e_ through time estimated from L0a carriers of separate Bantu speaker groups. Bayesian Skyline analysis for L0a carriers in North-Western Bantu speakers was not possible due to the low number of individuals with this haplogroup among them.

**Supplementary Figure 28:**
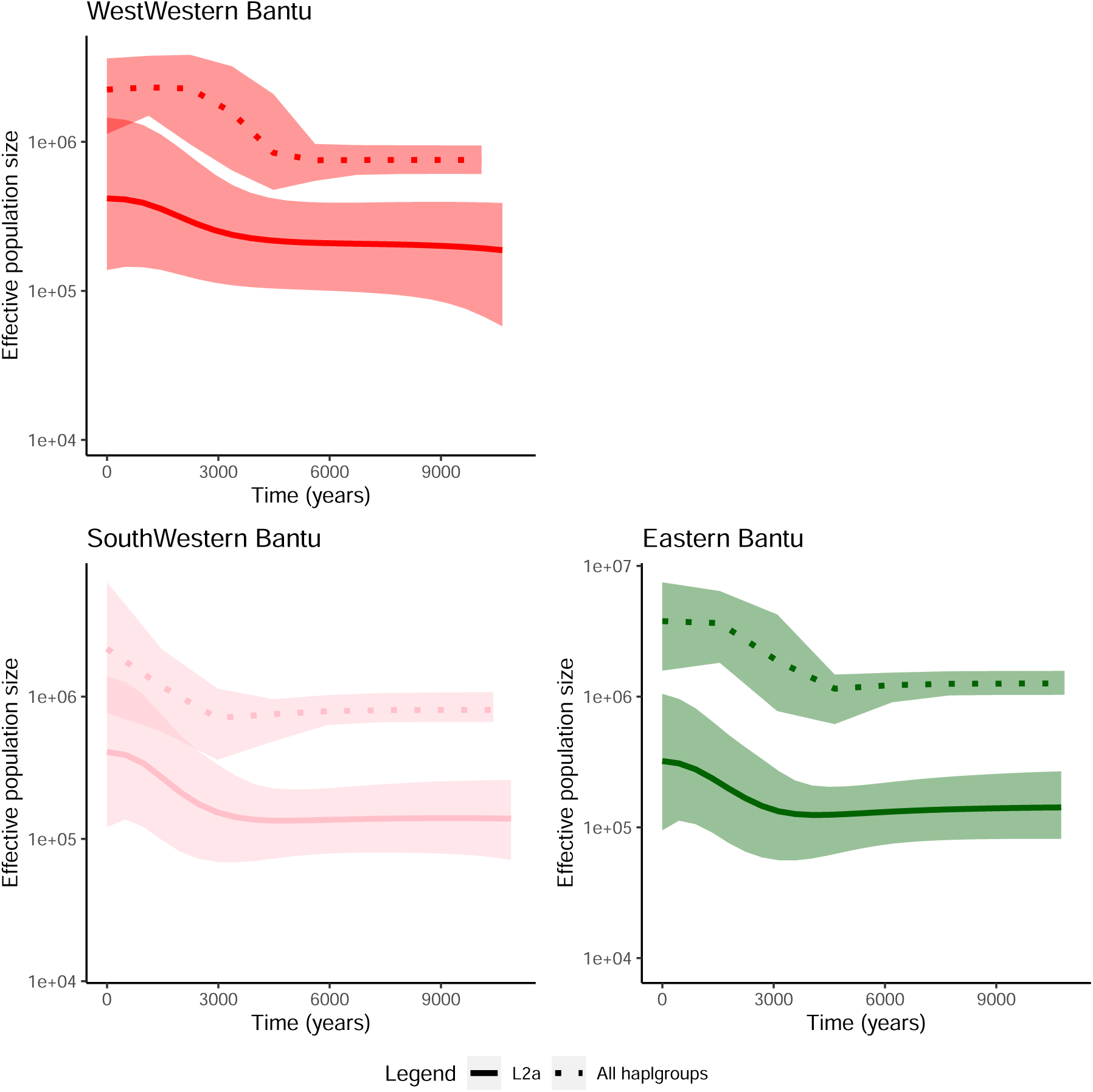
Bayesian Skyline Plots showing the variation of *N*_e_ through time estimated from L2a carriers of separate Bantu speaker groups. Bayesian Skyline analysis for L2a carriers in North-Western Bantu speakers was not possible due to the low number of individuals with this haplogroup among them.

**Supplementary Figure 29:**
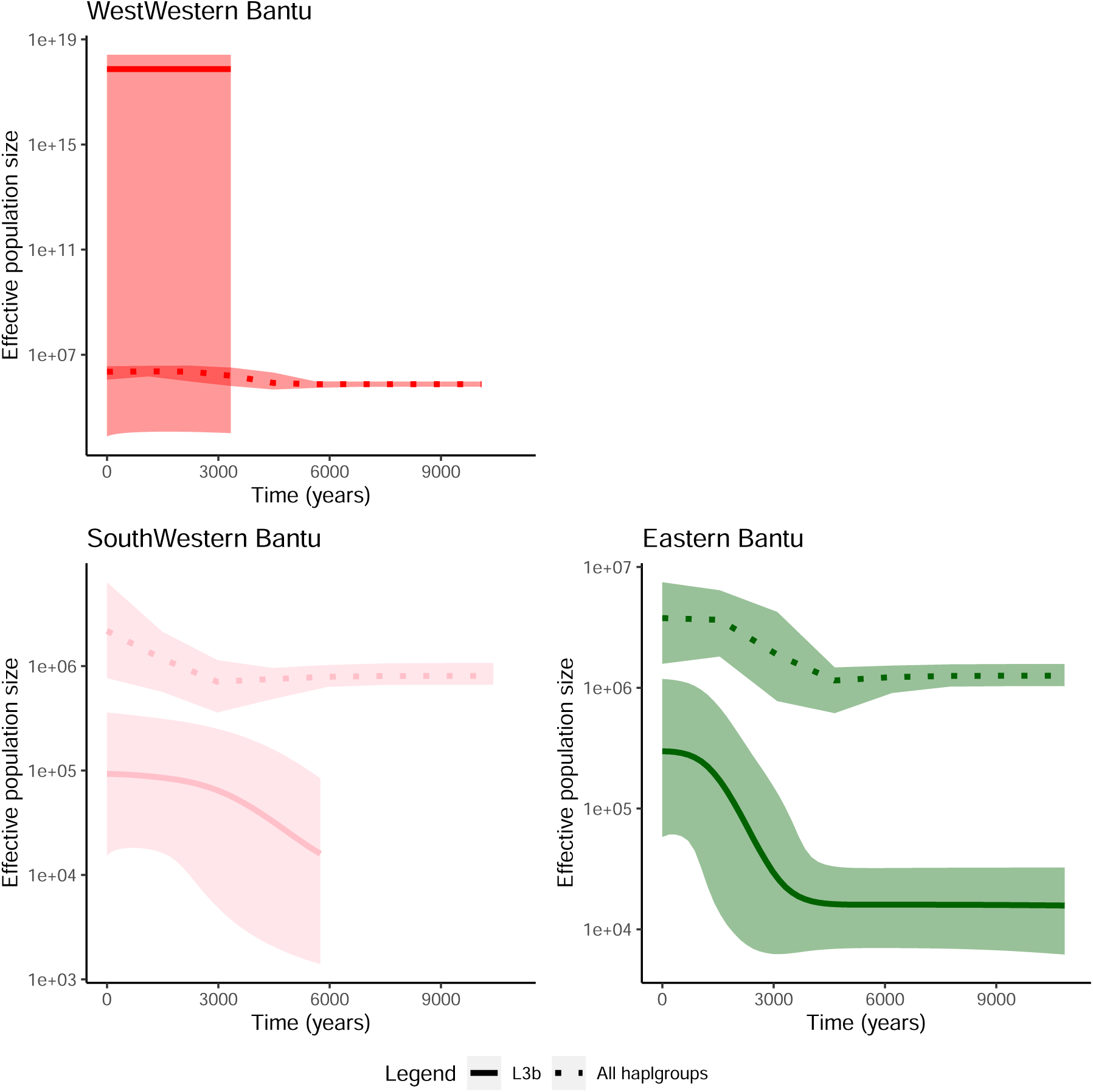
Bayesian Skyline Plots showing the variation of *N*_e_ through time estimated from L3b carriers of separate Bantu speaker groups. Bayesian Skyline analysis for L3b carriers in North-Western Bantu speakers was not possible due to the low number of individuals with this haplogroup among them. Due to the small number of West-Western Bantu speakers carrying haplogroup L3b (N=10), confidence intervals are large for the female *N*_e_ and should be interpreted with caution.

## Supplementary Tables

**Supplementary Table 1:** Ethical clearance and sample permission details for newly generated study samples. Ethical clearance and sample permission information for the newly generated samples in this study, including associated countries, Swedish reference numbers, local reference numbers, and the corresponding local institutions.

| Collection name | Country | Swedish reference number | Local reference number | Local institution |
| --- | --- | --- | --- | --- |
| DRC samples | DRC | Dnr 2019-05244 and Dnr 2021-01448 | Nr 091/CAB/MIN/-CA/PKB/2018 | Minister of Arts and Culture of the DRC |
| Ethiopian samples | Ethiopia | Dnr 2021-01448 | 310/169/2018 | The Federal Democratic Republic of Ethiopia Ministry of Science and Technology |
| Khoe-San and Bantu samples from Africa | Zimbabwe, South Africa, Namibia, Uganda, Zambia, Zanzibar | Dnr 2021-01448 | M180654, M180655, M180656 | Human Research Ethics Committee from Witwatersrand University |
| Khoe-San and Bantu samples from Botswana | Botswana | Dnr 2019-00115 | Permission obtained from the Botswana government in accordance with national regulations at the time of sampling | The Botswana Governmental Office |
| Sahel samples | Chad, Sudan, Senegal | Dnr 2021-01448 | 2016-07 | Institutional research board of Charles University, Prague |
| South African samples | South Africa | Dnr 2021-01448 | EC160429-024 (Health Ethics reference 259/2016) | Faculty of Natural and Agricultural Sciences University of Pretoria |
| Cameroon samples | Cameroon | Not applicable | CBI/397/ERCC/CAMBIN | Ethics Review and Consultancy Committee (ERCC), Cameroon Bioethics Initiative (CAMBIN) |
| Coloured, Khoe-San and Khoe-San descendants | South Africa | Dnr 2021-01448 | M980553 (renewals M050902, M090576, M1604104) | Human Research Ethics Committee from Witwatersrand University |
| Togo samples | Togo | Dnr 2021-04382 | N097/MESR/SG/-DRST/19 | Ministry of higher education and research |

**Supplementary Table 2:**
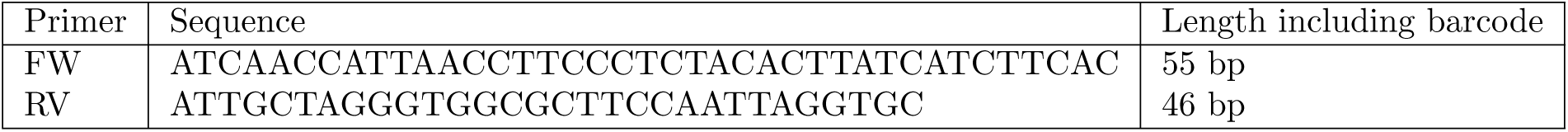
Primer sequences used for full mtDNA amplification using long-range PCR. The length of the primers, including the barcode is given in the last column.

**Supplementary Table 3:**
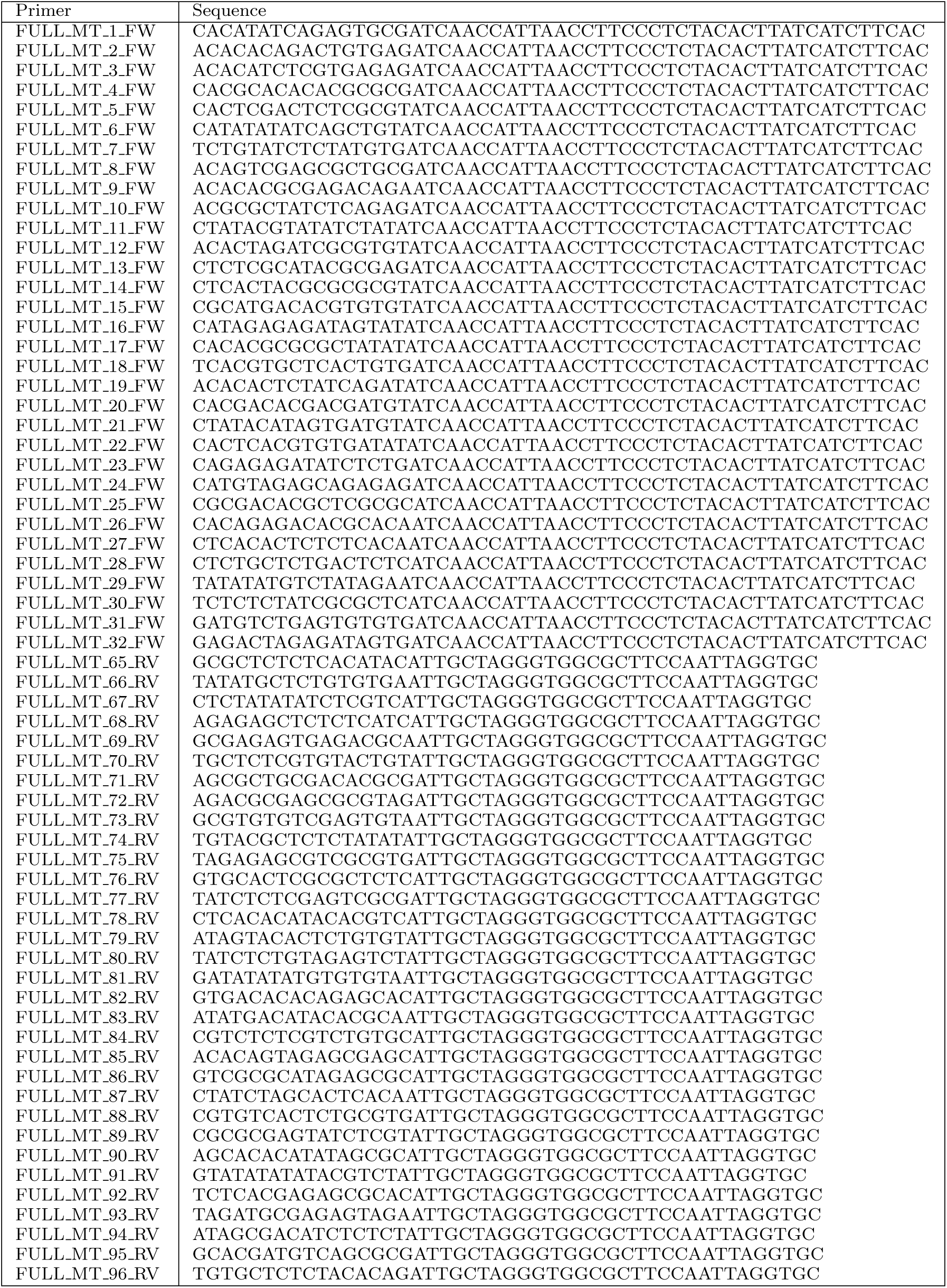
Barcoded primers used for amplification of the mtDNA sequences. The number of forward (32) and reverse (32) primers could create 1,024 unique combinations.

**Supplementary Table 4.** Ancestry associated by literature with the mitochondrial haplogroups. References are given in the last column.

| Haplogroup | Associated ancestry | Reference |
| --- | --- | --- |
| B4 | Asian | 104 |
| B5a | Asian | 104 |
| F1a | Asian | 104 |
| M | Asian | 104 |
| R | Asian | 104 |
| R0 | Asian | 104 |
| R5 | Asian | 104 |
| R6 | Asian | 104 |
| U2b | Asian | 104 |
| U2c | Asian | 104 |
| U2d | Asian | 105 |
| U7 | Asian | 104 |
| L0a | West-Central Africa (Bantu) | 52,53,106 |
| L1' | West-Central Africa (Bantu) | 107 |
| L1a | West-Central Africa (Bantu) | 54 |
| L1b | West-/Central Africa (Bantu) | 108 |
| L1c | West-Central Africa (Bantu) | 106 |
| L1e | West-Central Africa (Bantu) | 58,59 |
| L2 | West-Central Africa (Bantu) | 58,59 |
| L2' | West-Central Africa (Bantu) | 107 |
| L2a | West-Central Africa (Bantu) | 106 |
| L2b | West-Central Africa (Bantu) | 62 |
| L2c | West-Central Africa (Bantu) | 62 |
| L2d | West-Central Africa (Bantu) | 62 |
| L2e | West-Central Africa (Bantu) | 61 |
| L3b | West-Central Africa (Bantu) | 57 |
| L3d | West-Central Africa (Bantu) | 62 |
| L3e | West-Central Africa (Bantu) | 58,59,106,54 |
| L0b | East African | 51 |
| L0f | East African | 109 |
| L3a | East African | 109 |
| L3c | East African | 67 |
| L3f | East African | 109 |
| L3h | East African | 109 |
| L3i | East African | 109 |
| L3x | East African | 109 |
| L4 | East African | 109 |
| L4 | East African | 110 |
| L4a | East African | 110 |
| L4b | East African | 109 |
| L5 | East African | 109 |
| L5a | East African | 109 |
| L5b | East African | 109 |

| Haplogroup | Associated ancestry | Reference |
| --- | --- | --- |
| L6a | East African | 110 |
| L6b | East African | 110 |
| M1a | East African | 109 |
| M1b | East African | 111 |
| R0a | East African | 112 |
| B4a | East Asian | 104 |
| B4b | East Asian | 104 |
| B5b | East Asian | 113 |
| H | European | 104 |
| H1a | European | 114 |
| H1c | European | 114 |
| H5a | European | 114 |
| H53 | European | 3 |
| HV | European | 104 |
| HV+ | European | 104 |
| HV1 | European | 104 |
| I2' | European | 3 |
| J1c | European | 115 |
| J1d | European | 3 |
| J2a | European | 3 |
| K | European | 104 |
| K1a | European | 116 |
| N | European | 3 |
| N1a | European | 3 |
| N1b | European | 3 |
| T2b | European | 104 |
| U2e | European | 104 |
| U3a | European | 3 |
| U4' | European | 3 |
| U5a1 | European | 104 |
| U8b | European | 117 |
| V | European | 3 |
| L0d | KhoeSan | 1,50,8 |
| L0k | KhoeSan | 1,50,8 |
| H1v | North African | 118 |
| L3k | North African | 107 |
| U6a | North African | 119 |
| U6b | North African | 119 |
| U6c | North African | 119 |
| U6d | North African | 119 |
| L0 | Other | 1 |
| L3 | Other | 107 |
| L3' | Other | 107 |

| Haplogroup | Associated ancestry | Reference |
| --- | --- | --- |
| E1a | South Asian | 120 |
| M18 | South Asian | 121 |
| M2a | South Asian | 121 |
| M2b | South Asian | 121 |
| M33 | South Asian | 121 |
| M42 | South Asian | 122 |
| M5a | South Asian | 121 |
| M6a | South Asian | 3 |
| U2a | South Asian | 123 |
| U7a | South Asian | 123 |
| L0g | Unknown | No reference |

**Supplementary Table 5:** Settings for BEAST analyses by group: number of individuals, MCMC iterations, replicates, burn-in, and total iterations used for for Bayesian phylogenies. The table presents the configurations used for BEAST (Bayesian Evolutionary Analysis Sampling Trees) analyses, organized by language groups. Each group is characterized by the number of individuals within it. The BEAST analyses entail multiple runs, each involving a specific number of Markov Chain Monte Carlo (MCMC) iterations, conducted in replicate. These replicates are subsequently merged to enhance the accuracy of Bayesian phylogeny inferences. The table also notes the ‘burn-in’ period discarded during the analysis and the total number of iterations employed in generating the Bayesian Skyline Plot.

| Group<br>(language/haplogroup) | Number<br>of individuals | Iterations<br>(million) | Replicate runs<br>performed | Discarded<br>burn-in (million) | Total iterations used<br>for BSP (million) |
| --- | --- | --- | --- | --- | --- |
| Nilo-Saharan | 81 | 10 | 20 | 0.06 | 198.8 |
| Afrikaans | 211 | 100 | 10 | 1 | 990 |
| Afrikaans, L0d and L0k | 126 | 100 | 10 | 10 | 900 |
| Afrikaans, Indo-European haplogroups | 18 | 100 | 10 | 10 | 900 |
| Afro-Asiatic | 396 | 100 | 10 | 2 | 980 |
| Khoisan | 783 | 100 | 20 | 2 | 1960 |
| Niger-Congo (non Bantu) and Mande | 844 | 20 | 58 | 1.5 | 647.5 |
| Bantu | 1746 | 15 | 60 | 2 | 180 |
| North-Western Bantu | 30 | 10 | 20 | 0.03 | 199.4 |
| Eastern Bantu | 609 | 20 | 50 | 1.5 | 925 |
| West-Western Bantu | 320 | 100 | 20 | 0.4 | 1992 |
| South-Western Bantu | 787 | 50 | 19 | 0.7 | 936.7 |
| L0d | 856 | 10 | 20 | 1 | 720 |
| L0k | 120 | 10 | 10 | 0.2 | 98 |
| L3e Eastern Bantu | 89 | 100 | 10 | 0.1 | 999 |
| L3e West-Western Bantu | 71 | 100 | 10 | 0.1 | 999 |
| L3e North-Western Bantu | 7 | 100 | 10 | 0.1 | 999 |
| L3e South-Western Bantu | 154 | 100 | 10 | 0.1 | 999 |
| L1c North-Western Bantu | 18 | 100 | 10 | 1 | 990 |
| L1c West-Western Bantu | 58 | 100 | 10 | 1 | 990 |
| L1c South-Western Bantu | 131 | 100 | 10 | 1 | 990 |
| L1c Eastern Bantu | 62 | 100 | 10 | 1 | 990 |
| L0a West-Western Bantu | 37 | 100 | 10 | 1 | 990 |
| L0a South-Western Bantu | 104 | 100 | 10 | 1 | 990 |
| L0a Eastern Bantu | 97 | 100 | 10 | 1 | 990 |
| L2a West-Western Bantu | 46 | 100 | 10 | 1 | 990 |
| L2a South-Western Bantu | 86 | 100 | 10 | 1 | 990 |
| L2a Eastern Bantu | 75 | 100 | 10 | 1 | 990 |
| L3b South-Western Bantu | 13 | 100 | 10 | 1 | 990 |
| L3b Eastern Bantu | 43 | 100 | 10 | 1 | 990 |

**Supplementary Table 6:** Time to the most recent common ancestor (TMRCA) of all individuals belonging to certain groups.

| Group | TMRCA (kya) | TMRCA 95% confidence interval (kya) |
| --- | --- | --- |
| Modern humans and Neanderthals | 304.7 | 267.0 - 344.1 |
| All modern humans | 132.4 | 117.4 - 147.8 |
| L0d | 116.3 | 91.9 - 142.8 |
| L0f | 82.5 | 69.3 - 95.3 |
| L0k | 64.7 | 47.5 - 82.8 |
| L0a | 67.4 | 54.2 - 81.6 |
| L1c | 81.7 | 70.6 - 93.2 |
| L2a | 58.4 | 48.5 - 69.1 |
| L3e | 36.5 | 30.2 - 43.4 |

## Notes

### Competing Interest Statement

The authors have declared no competing interest.

